# Cholesterol differentially modulates the activity of opioid and muscarinic receptors via a common binding site

**DOI:** 10.1101/2024.10.09.617372

**Authors:** Nikolai Chetverikov, Alena Janoušková-Randáková, Dominik Nelic, Jan Jakubík

## Abstract

G protein-coupled receptors (GPCRs) are membrane proteins that represent the largest and most therapeutically targeted receptor class, accounting for 30% of currently marketed drugs. Two binding motifs for membrane cholesterol, the cholesterol recognition amino acid consensus (CRAC) domain and the cholesterol consensus motif (CCM), have been postulated. Using a simulation of the molecular dynamics of cholesterol association with the receptor, we predicted the binding of membrane cholesterol to non-canonical sites, distinct from CRAC and CCM, at muscarinic and opioid receptors. We identified a binding site common to muscarinic and opioid receptors at TM6, with arginine 6.35 as the major residue. Membrane cholesterol depletion mimics the effects of R^6.35^ mutations, confirming its role in receptor modulation. Targeting cholesterol-binding sites offers novel pharmacotherapeutic strategies, including tissue-specific sterol-based modulation.

**One sentence summary:** This study identifies a shared non-canonical cholesterol-binding site at TM6 in muscarinic and opioid receptors, with significant implications for drug development targeting tissue-specific GPCR modulation.

## 1 Introduction

G protein-coupled receptors (GPCRs) are receptors that mediate diverse physiological functions. Over 800 members of the GPCR superfamily comprise the largest and most therapeutically targeted receptor class, representing 30% of currently marketed drugs[1]. Numerous studies have shown that cholesterol interacts with GPCRs at specific sites, directly allosterically modulating ligand binding and affecting their activation and pharmacology[2]. Muscarinic acetylcholine receptors and opioid receptors are integral components of the central nervous system (CNS), playing crucial roles in regulating various physiological and neurological functions. Both receptor types are involved in mediating essential processes such as cognition, mood regulation, pain perception, and autonomic functions. Understanding their mechanisms and implications in health and disease is vital for developing targeted therapies.[3].

Muscarinic receptors are a subtype of acetylcholine receptors that are member of G protein-coupled receptors (GPCRs) family. They are predominantly found in the CNS and peripheral tissues. In the brain, these receptors influence memory formation, higher cognitive functions, mood, and behavioural responses[4]. Malfunction or dysregulation of muscarinic receptors has been linked to several neurological and psychiatric conditions, including dementias such as Alzheimer’s disease, schizophrenia, anxiety disorders, and depression. Thus, muscarinic receptors emerged as validated targets for the symptomatic treatment of these conditions[5]. Opioid receptors, also GPCRs, are primarily involved in modulating pain, mood, and sedation. They mediate analgesic effects, which are crucial in pain management, but also influence emotional states, contributing to feelings of depression or dysphoria. Additionally, opioid receptors are implicated in sleep regulation. Due to their widespread influence, they are targeted pharmacologically for treating acute and chronic pain, major depressive disorder, and sleep disturbances.[6].

Beyond their central roles, both muscarinic and opioid receptors are present in peripheral tissues, where they regulate functions such as glandular secretion, gastrointestinal motility, and cardiovascular responses [4]. Activation of these receptors with agonists can lead to adverse side effects. For instance, muscarinic agonists may cause excessive sweating and incontinence, while opioid agonists can induce constipation and respiratory depression. These side effects pose significant challenges in clinical applications.To mitigate adverse effects, researchers are exploring alternative approaches such as the use of positive allosteric modulators (PAMs). PAMs enhance the signalling of endogenous ligands without directly activating the receptor, thereby maintaining the natural spatiotemporal signalling patterns. This approach offers increased selectivity and potentially fewer side effects compared to traditional orthosteric ligands. The development of PAMs represents a promising avenue for creating more effective and safer therapeutic agents targeting muscarinic and opioid receptors.[7–10].

Cholesterol plays a crucial role in the structure and function of various membrane proteins, particularly G protein-coupled receptors (GPCRs). It has been observed to co-crystallize with numerous GPCRs, indicating direct interactions that influence receptor behaviour and signalling pathways[11]. Cholesterol binding to GPCRs affects ligand binding and functional response as well as receptor oligomerization and trafficking [12–14]. Researchers have proposed two primary binding motifs for membrane cholesterol, which are essential for understanding these interactions. The first is the well-known ‘Cholesterol Recognition Amino Acid Consensus’ (CRAC) domain, characterized by the pattern (–L/V-(X)1–5-Y-(X)1–5-R/K-), found across many membrane proteins. This motif facilitates specific binding of cholesterol through its amino acid sequence, contributing to the stability and function of the protein within the lipid bilayer[15]. Subsequently, a (K/R)-X1−5-(Y/F)-X1−5-(L/V) reversed pattern (CARC) was found in the nicotinic acetylcholine receptor[16]. Mutagenesis of CRAC/CARC residues reduced cholesterol binding or cholesterol-dependent function (e.g. peripheral benzodiazepine receptor (TSPO)[17], nicotinic acetylcholine receptor[18], voltage-gated ion channels[19], GPCRs[20] and other membrane proteins), confirming this motif experimentally. However, not all predicted CRAC or CARC motifs bind cholesterol effectively, as their binding capacity is highly context-dependent. Factors such as the orientation of the motif within the membrane, lipid exposure, and the surrounding structural environment significantly influence binding affinity and specificity[21]. Beyond linear motifs, a three-dimensional ‘Cholesterol Consensus Motif’ (CCM) has been identified through structural studies, notably in the β_2_-adrenergic receptor (structure ID: 3D4S). This motif suggests that cholesterol binding can also occur via spatial arrangements within the receptor’s transmembrane domains, and similar motifs are predicted in other GPCRs, including muscarinic acetylcholine receptors[22].

While certain motifs like the Cholesterol Recognition Amino-acid Consensus (CRAC) sequence are known to facilitate cholesterol interactions with some GPCRs, not all GPCRs possess these motifs. While canonical motifs like CRAC are well-characterized, some cholesterol interactions occur through non-traditional mechanisms. Certain α-helical peptides can bind cholesterol without possessing the classic basic or aromatic residues, forming ‘tilted’ binding domains. These domains accommodate cholesterol by tilting their hydrophobic face into the membrane, creating topological sites for sterol binding[21]. Additionally, structural studies using X-ray crystallography often reveal cholesterol molecules bound in non-annular sites—positions located between transmembrane helices of GPCRs or ion channels—highlighting the complexity and diversity of cholesterol-protein interactions in the membrane environment[23].

Although many GPCRs lack “canonical” cholesterol-binding sites, cholesterol still modulates their function, indicating that specific interactions beyond traditional motifs are at play[24, 25]. A current extensive study of cryo-EM and X-ray structures has shown that almost all cholesterol binding to GPCRs occurs in predictable locations that, however, lack perceivable cholesterol-binding motifs[26]. Cholesterol can enhance the conformational stability of GPCRs, either by promoting active or inactive states, which is crucial for their signalling functions. This stabilisation is often achieved through direct interactions with specific residues rather than relying solely on conserved motifs[11, 24]. The high plasticity of cholesterol interaction sites and the lack of universal motifs highlight the importance of detailed, residue-level analysis to understand how cholesterol influences GPCR function. This is supported by studies that use molecular dynamics simulations and structural bioinformatics to elucidate cholesterol-GPCR interactions at a finer scale[11, 25].

While “canonical” cholesterol-binding motifs like CRAC and CCM have been extensively studied, “non-canonical” binding sites remain poorly understood. This study identifies and characterises a shared “non-canonical” site at TM6 in muscarinic and opioid receptors. In our previous study, we have shown that membrane cholesterol prevents persistent activation of muscarinic receptors by wash-resistant xanomeline by interaction with R^6.35^ and L/I^6.46^(Ballesteros and Weinstein numbering[27]) in the TM6[28]. The identified cholesterol-binding site does not correspond to any known cholesterol-binding motif. Similarly to M_2_ and M_4_ muscarinic receptors, a substantial number of G_i/o_-coupled class A GPCRs possess a basic residue in position 6.35 and hydrophobic L/I/V in position 6.46 next to a conserved proline kink (Figure 1). We speculate that membrane cholesterol also binds to R^6.35^ at opioid receptors. In the present study, we docked cholesterol to M_2_ and M_4_ muscarinic receptors and δ-, κ- and μ-opioid receptors by simulation of molecular dynamics of association of cholesterol with receptors[29]. We identified two common cholesterol-binding sites, including the one containing R^6.35^. Then we show that depletion of membrane cholesterol has similar effects on functional responses of tested receptors as mutations of R^6.35^ aimed at disruption of cholesterol binding to this residue, confirming a common non-canonical binding site at R^6.35^. Variation in interactions of receptors with the membrane represents a possibility to achieve pharmacological selectivity based on variation in membrane composition among tissues, affording tissue-specific selectivity.

**Figure 1.**
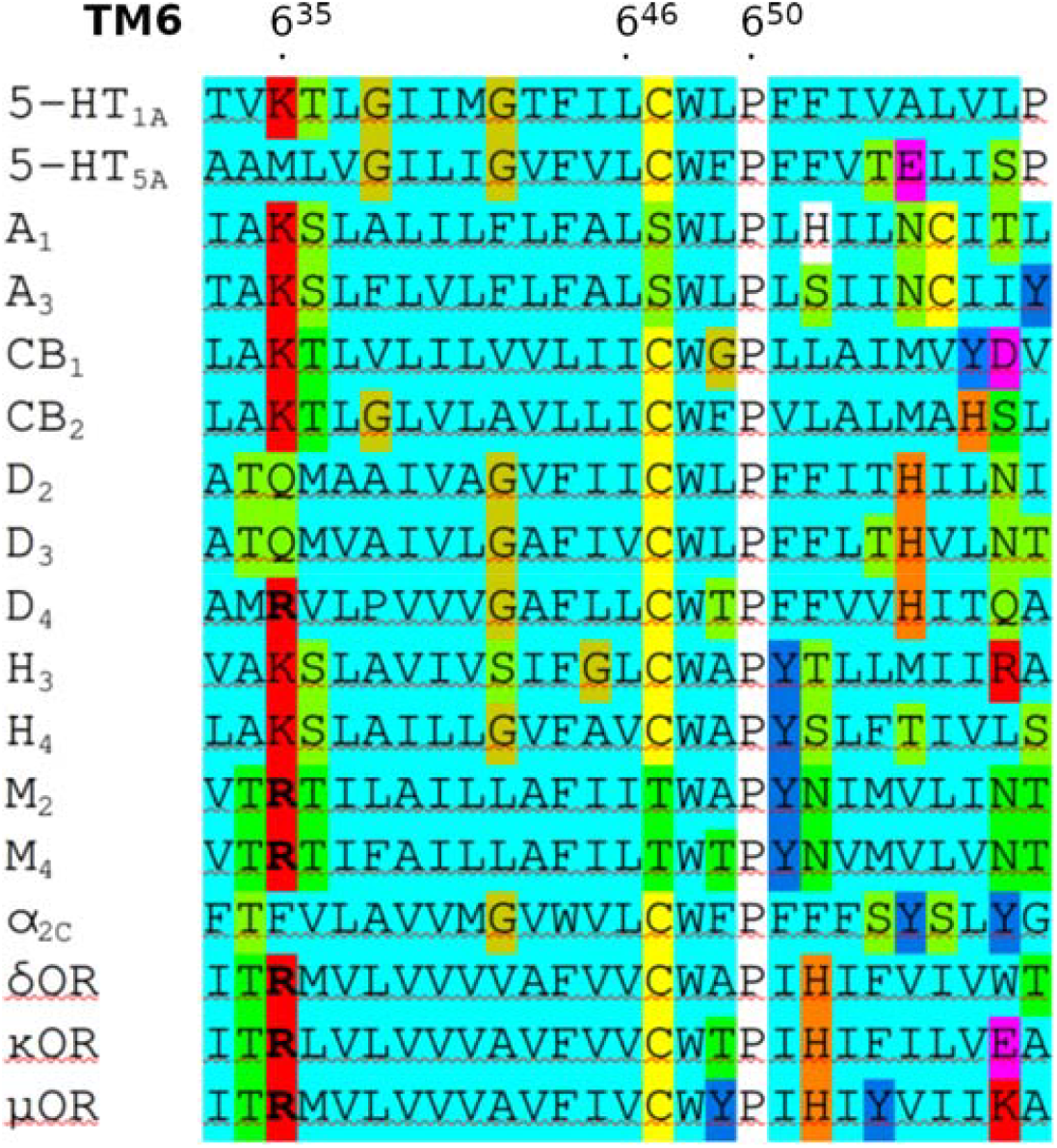
Protein sequences of TM6. Aligned protein sequences of the sixth transmembrane α-helix (TM6) of selected G_i/o_ coupled GPCRs of class A. Cyan, non-polar; green, polar; blue, aromatic polar; yellow, cysteine; gold, glycine; white, proline; magenta, acidic; red, basic amino acids. Bold, arginine.

## 2 Materials and Methods

### 2.1 Materials

Common chemicals were purchased from Sigma (Prague, CZ) in the highest available purity. Radiolabelled [^35^S]GTPγS, [^3^H]DAMGO, and [^3^H]diprenophin were Perkin Elmer NEG030H, NET902, and NET11212, respectively, and [^3^H]N-methyl scopolamine was American Radiochemical Company ART1214 purchased from Lacomed (Kralupy nad Vltavou, CZ).

### 2.2 Cell culture and membrane preparation

We transfected Chinese hamster ovary (CHO) cells with the genes of individual human variants of muscarinic and opioid receptors (Missouri S&T cDNA Resource Center, Rolla, MO, USA). CHO cell cultures were prepared as described previously. Stable cell lines were grown to confluence in 75 cm^2^ flasks in Dulbecco’s modified Eagle’s medium supplemented with 10% fetal bovine serum, and 2 million cells were subcultured in 100 mm Petri dishes. The medium was supplemented with 5 mM butyrate for the last 24 hours of cultivation to increase receptor expression. Cells were detached by mild trypsinisation on day 5 after subculture. For the cultivation of transiently transfected cells, see “Construction of mutant receptors” below. Detached cells were washed twice in 50 ml of phosphate-buffered saline and 3 min centrifugation at 250 x g. Harvested cells were stored at −20°C overnight. Thawed cells were suspended in 20 ml of ice-cold incubation medium (100 mM NaCl, 20 mM Na-HEPES, 10 mM −MgCl_2_, pHCC7.4) supplemented with 10 mM EDTA and homogenised on ice by two 30-s strokes using a Polytron homogeniser (Ultra-Turrax; Janke & Kunkel GmbH & Co. KG, IKA-Labortechnik, Staufen, Germany) with a 30-s pause between strokes. Cell homogenates were centrifuged for 5 min at 1000×g to remove whole cells and cell nuclei. The resulting supernatants were centrifuged for 30 min at 30,000×g. Pellets were suspended in a fresh incubation medium, incubated on ice for 30 min, and centrifuged again. The resulting membrane pellets were kept at −80 °C until assayed within 10 weeks.

### 2.3 Depletion of membrane cholesterol

We reduced membrane cholesterol levels by incubating whole detached cells in Krebs-HEPES buffer (KHB; final concentrations in mM: NaCl 138; KCl 4; CaCl_2_ 1.3; MgCl_2_ 1; NaH_2_PO_4_ 1.2; Hepes 20; glucose 10; pH adjusted to 7.4) supplemented with 10 mM methyl-β-cyclodextrin (MβCD) at 37ºC for 30 min. The concentration of the cells in suspension was 20 million cells per ml. After incubation, the medium was removed, and cells were washed twice with a triple incubation volume of fresh KHB. Effects of treatments on cholesterol levels were determined using Amplex® Red Cholesterol Assay Kit (Molecular Probes, Eugene, OR) according to the manufacturer’s protocol.

### 2.4 Construction of mutant receptors

Plasmids coding muscarinic and opioid receptors were obtained from the Missouri S&T cDNA Resource Center (Rolla, MO, USA) and used to generate sequences with the desired amino acid substitution using Q5 Site-Directed Mutagenesis Kit (NEB, via BioTek, Praha, Czech Republic). Primers were designed using NEBaseChanger (https://nebasechanger.neb.com/). CHO-K1 cells were transfected with desired plasmids using linear polyethylenimine (PEI) (Polysciences, Germany). Approximately 2 million cells were seeded in 10 ml of a 1:1 mixture of Dulbecco’s modified Eagle’s and Ham’s F-12 with L-glutamine in a 100 mm Petri dish and incubated for 24 hours at 37°C in a humidified 5% CO_2_ atmosphere. DNA and PEI were mixed in a 1:3 ratio (12 μg DNA per 1 ml PBS : 36 μg PEI per 1 ml PBS) and incubated at room temperature for 30 min. Then 1 ml of transfection mix was added to a dish and incubated for 16 hours at 37°C in a humidified 5% CO_2_ atmosphere. Then DMEM was removed, and 8 ml of fresh DMEM was added, and cells were incubated for an additional 24 hours.

### 2.5 [^35^ S]GTPγS binding

Measurement of [^35^S]GTPγS binding was carried out on membranes in 96-well plates at 30°C in the incubation medium described above supplemented with freshly prepared dithiothreitol at a final concentration of 1 mM, essentially as described by [30]. Membranes at concentrations of 10 μg of protein per well were used. The final volume was 200 μl, the final concentration of [^35^S]GTPγS was 500 pM, and the final concentration of GDP was 50 μM. Non-specific binding was determined in the presence of 1 μM unlabelled GTPγS. Total GTP binding capacity was determined in the absence of GDP. Agonist was added 15 min before [^35^S]GTPγS. Incubation with [^35^S]GTPγS was carried out for 20 min, and the free ligand was removed by filtration through GF/C glass-fibre filtration plates (Millipore or Unifilter) using a Brandel cell harvester (Brandel, Gaithersburg, MD, USA). Filtration and washing with ice-cold water lasted for 9 seconds. Filters were dried, and then 40 µl of liquid scintillator Rotiszint was added to each well. The filtration plates were counted in a Wallac-Microbeta scintillation counter.

### 2.6 Saturation binding

Membranes (20-50 μg of membrane proteins per sample) were incubated in 96-well plates for 3 h at 30°C in 800 μl (200 μl for κ-OR) of incubation medium with [^3^H]deltrophin (δ-OR), [^3^H]diprenorphine (κ-OR), [^3^H]DAMGO (μ-OR) or [^3^H]N-methylscopolamine ([^3^H]NMS, M and M) at six concentrations of radioligand in two-fold dilutions. Nonspecific binding was determined in the presence of 10 μM Leu-enkephalin (opioid receptors) or 1 μM atropine (muscarinic receptors). Incubations were terminated by filtration through GF/C glass-fibre filtration plates (Millipore or Unifilter) using a Brandel cell harvester (Brandel, Gaithersburg, MD, USA). Filtration and washing with ice-cold water lasted for 6 seconds. Filters were dried, and then 40 µl of liquid scintillator Rotiszint was added to each well. The filtration plates were counted in a Wallac-Microbeta scintillation counter.

### 2.7 Experimental data analysis

Experiments were independent. Functional assays were carried out in quadruplicate. After subtraction of non-specific binding, data were normalised to the total GTP binding capacity determined in each experiment. EC_50_ values were calculated as logarithms.

Parameters of a functional response were determined by fitting Eq. 1 to experimental data.

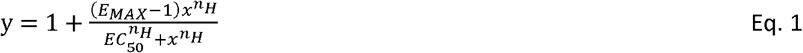

Where y is the functional response at a concentration of tested compound x, E_MAX_ is the maximal response to the tested compound, EC_50_ is the half-efficient concentration, and n_H_ is the slope factor (Hill coefficient).

Saturation binding experiments were carried out in triplicates. After subtraction of non-specific binding, data were converted to substance amounts and concentrations. Eq. 2 was fitted to data from saturation experiments.

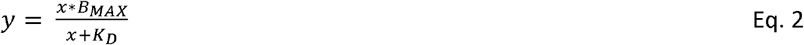

Where y is the specific radioligand binding at a concentration of the radioligand x, B_MAX_ is the maximal binding capacity, and K_D_ is the equilibrium dissociation constant of the radioligand.

### 2.8 Molecular modelling

#### 2.8.1 Preparation of receptor structure

Structures of the M_2_ and M_4_ muscarinic acetylcholine (3UON, 4MQS, 5DSG, 8FX5) and μ, κ and δ opioid (4DKL, 5C1M, 4DJH, 6B73, 4EJ4, 6PT2) and A_2A_ adenosine (5IU4) and β_2_ adrenergic (3D4S) receptors were downloaded from RCSB Protein Data Bank (https://www.rcsb.org/). Non-protein molecules (except ligands) and nanobody or hydrolase parts of protein were deleted, and the resulting receptor protein was processed in Maestro using Protein Preparation Wizard according to Sastry et al. guidelines[31]. No missing receptor parts (e.g. the third intracellular loop, N-terminus, C-terminus) were constructed.

#### 2.8.2 Coarse-grained MD

The coarse-grained molecular dynamics (cgMD) was used to simulate the association of cholesterol with GPCRs[44]. Pre-processed receptor molecules were uploaded to the CHARMM-GUI server (charmm-gui.org)[45]. Then, the Martini force field 3.0[46] coarse-grain system consisting of the receptor immersed in a 20 nm x 20 nm patch of POPC (1-palmitoyl-2-oleoyl-sn-glycero-3-phosphocholine) membrane containing 30 % of cholesterol was created using Membrane-Builder[47] and Martini maker[48]. After minimisation and equilibration of the system, 3 independent runs of 2 µs molecular dynamics using Gromacs 2018 were run[49]. Input files for minimisation (10,000 steps, coulomb type reaction field and Verlet cutoff scheme), relaxation (250,000 20-fs steps, 300 K, velocity rescale temperature coupling, Berendsen pressure coupling at 1 bar, semi-isotropic coupling, coulomb type reaction field and Verlet cutoff scheme), and production (10^8^ 20-fs steps, 300 K, velocity rescale temperature coupling, Parrinello-Rahman pressure coupling at 1 bar, semi-isotropic coupling, coulomb type reaction field and Verlet cutoff scheme) runs were created by CHARMM-GUI.

#### 2.8.3 Cholesterol re-docking

Molecules of cholesterol that associated with a receptor during cgMD were back-mapped onto in the first step (2.8.1) pre-processed receptor structure[50]. The resulting complexes were minimised in YASARA (NOVA force field)[51] and the cholesterol molecule was re-docked using the YASARA implementation of AutoDock[52]. The binding site was defined as a 5 Å extension in all directions to the cholesterol molecule, and the AutoDock local search method was employed; 888 poses were generated. All resulting poses were re-scored in YASARA by energy-minimisation of ligand pose and AutoDock VINA’s local search, confined closely to the original ligand pose. All top docking poses are deposited at Mendeley Data repository (doi:10.17632/3vyt4fbv4r.1).

#### 2.8.4 Simulation of molecular dynamics

To infer cholesterol-receptor interactions, the conventional MD (cMD) of the top-scoring pose (one cholesterol molecule per simulation) of individual cholesterol-receptor complexes was simulated to evaluate cholesterol binding to the receptor and assess its interactions with the receptor and potential effects on ligand binding. Simulation of cMD was performed using Desmond/GPU v6.5[53]. The simulated system consisted of the receptor-ligand-cholesterol complex in the POPC membrane in water and 0.15 M NaCl. The system was first relaxed by the standard Desmond protocol for membrane proteins and then 3 independent replicas of 600 ns NPγT (Noose-Hover chain thermostat at 300 K, Martyna-Tobias-Klein barostat at 1.01325 bar, isotropic coupling, and Coulombic cutoff at 0.9 nm, OPLS_2005 force field). The quality of the molecular dynamics simulation was assessed by the Simulation Quality Analysis tools of Maestro and analysed by the Simulation Event Analysis tool. Ligand-receptor interactions were identified using the Simulation Interaction Diagram tool.

#### 2.8.5 Statistical analysis

Receptor binding and functional response measurements were carried out in three independent experiments performed in quadruplicates. Three independent replicas of MD simulations were performed. Results are represented as means ± SD. To assess differences in obtained values t-test was used. Bonferroni correction for multiple comparisons was applied when necessary.

## 3 Results

### 3.1 Coarse-grained molecular dynamics

The coarse-grained MD (cgMD) was used to simulate the association of cholesterol to identify potential cholesterol-binding sites at muscarinic and opioid receptors[44]. The cgMD of a 20 nm x 20 nm patch of membrane containing 30% of cholesterol molecules and one receptor was simulated. Throughout cgMD, cholesterol molecules associated with the receptor and remained bound to it for hundreds of nanoseconds. To verify the approach, structures of the β_2_-adrenergic receptor (3D4S) and A_2A_-adenosine receptor (5IU4) were included in the test set. Cholesterol in locations appearing in these crystal structures was observed, validating the method. Namely, binding to the cholesterol consensus motif (CCM) between TM2 and TM4 of the β_2_-adrenergic receptor and TM2-TM3-TM4 extracellular bundle and TM6 extracellular part of the A_2A_-adenosine receptor was observed. Subsequently, cgMD was run at δ-opioid (4EJ4), κ-opioid (4DJH), μ-opioid (4DKL), M_2_-muscarinic (3UON) and M_4_-muscarinic (5DSG) receptors in an inactive conformation and δ-opioid (6PT2), κ-opioid (6B73) and μ-opioid (5C1M), M_2_-muscarinic (4MQS) and M_4_-muscarinic (8FX5) receptors in an active conformation. We have observed up to five concurrently bound molecules of cholesterol to eight unique binding sites (Figure 2) as detailed below.

**Figure 2.**
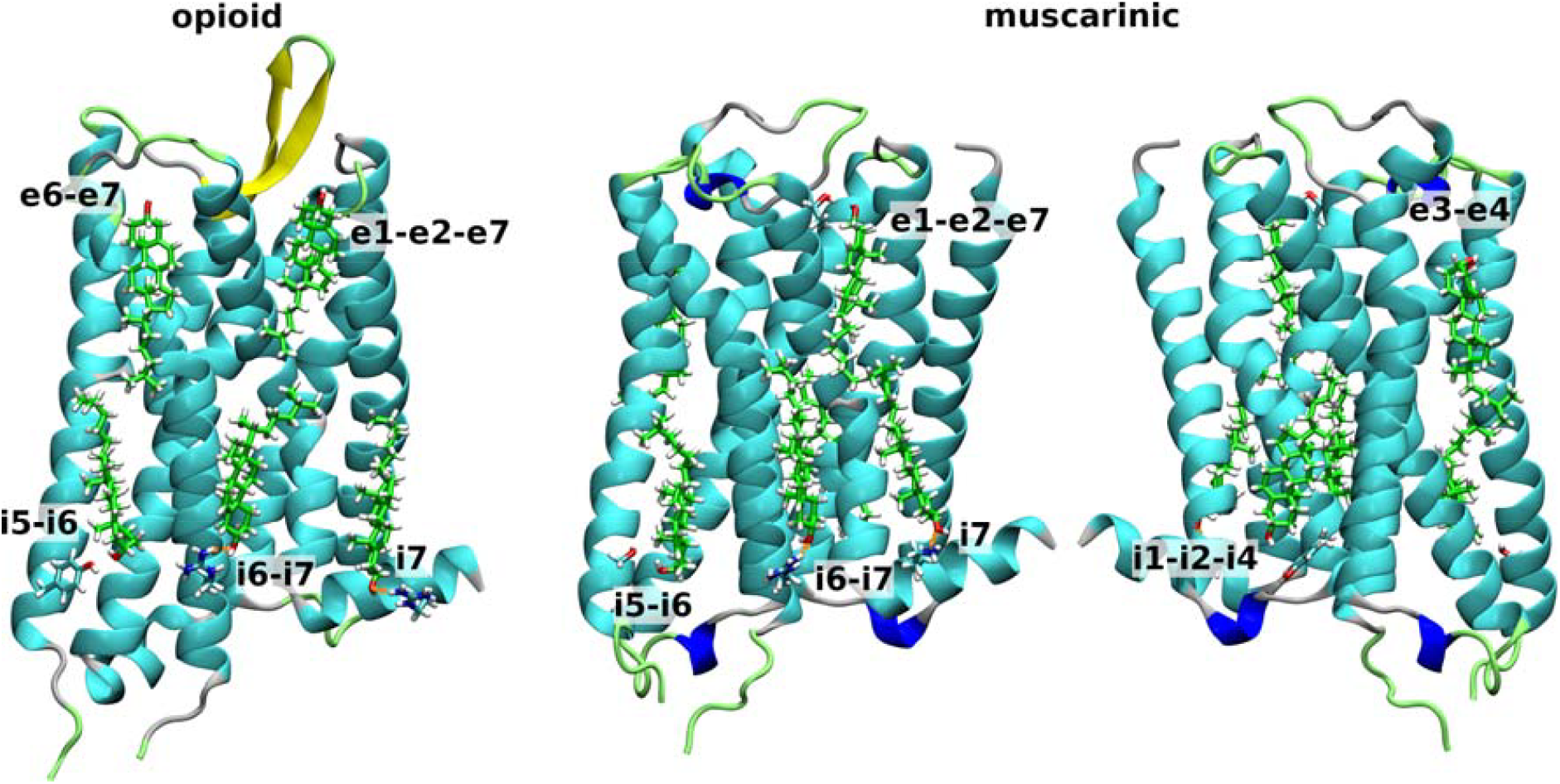
Cholesterol binding to opioid and muscarinic receptors. Molecules of cholesterol that associated with the opioid (left) or muscarinic receptor (middle and right) were back-mapped and re-docked to the receptor. Orientation: extracellular up, TM6 front (left and middle) or TM4 front (right). Cholesterol (green carbons) in the extracellular leaflet interacting with TM1-TM2-TM7 (e1e2-e7) bundle or TM3-TM4 (e3-e4) or TM6-TM7 (e6-e7) groove, and in the intracellular leaflet interacting with TM1-TM2-TM4 (i1-i2-i4) bundle or TM5-TM6 (i5-i6) or TM6-TM7 (i6-i7) groove or TM7 (i7) helix is shown.

### 3.2 Cholesterol re-docking

Eight unique cholesterol-binding sites on muscarinic and opioid receptors were identified using cgMD simulation of cholesterol association with the receptor, three in the extracellular and five in the intracellular leaflet of the membrane (Figure 2). Cholesterol molecules in the extracellular leaflet interacted with the TM1-TM2-TM7 (e1-e2-e7) bundle of helices or the groove between TM4-TM5 (e4-e5) or TM6-TM7 (e6/e7) helices. Cholesterol molecules in the intracellular leaflet interacted with the TM1-TM2-TM4 (i1-i2-i4) bundle of helices or the groove between TM3-TM4 (i3-i4), TM5-TM6 (i5-i6) or TM6-TM7 (i6-i7) helices or with TM7 (i7) helix. Specifically, cholesterol bound to four sites (e1-e2-e7, e6-e7, i3-i4 and i6-i7) at δ- and κ-opioid receptors, five sites (e1-e2-e7, e6-e7, i5-i6, i6-7 and i7) at the μ-opioid receptor (Figure 3) and six sites (e1-e2-e7, e4-e5, i3-i4, i5-i6, i6-i7 and i7) at muscarinic receptors (Figure 4). Individual molecules of cholesterol were back-mapped onto the receptor[50], re-docked[54] and rescored[55]. Re-docking binding energy estimates were above 4 kcal/mol. Two binding sites were common for all studied receptors: e1-e2-e7 and i6-i7. Hydrogen bonding to R^6.35^ was common to all receptors in at least one conformation (Figure 5). Binding to the e1-e2-e7 sites markedly varied among receptors.

**Figure 3.**
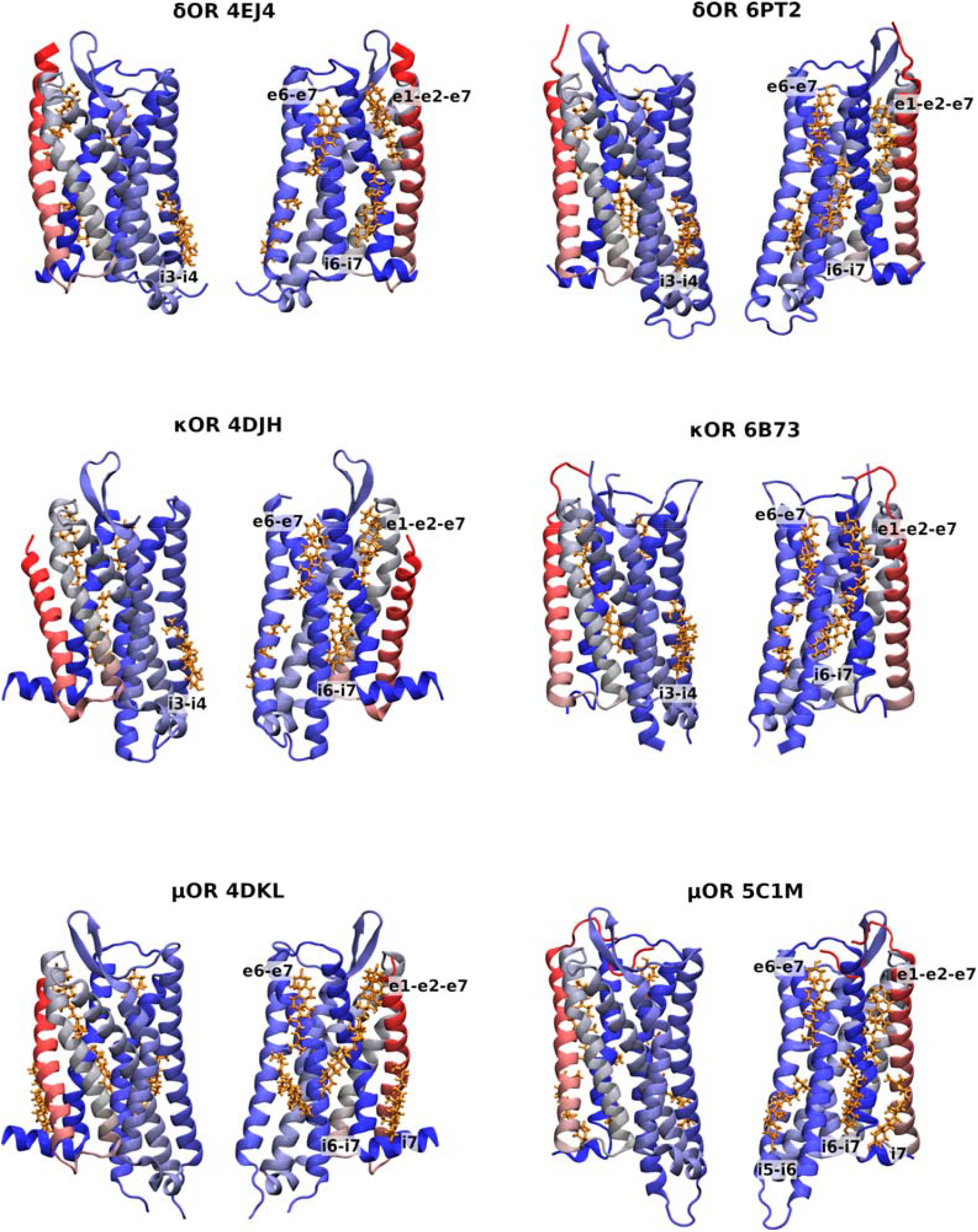
Cholesterol binding to opioid receptors. Molecules of cholesterol that are associated with opioid and muscarinic receptors in an inactive or active conformation during cgMD were back-mapped and re-docked to the receptors. Cholesterol is gold. The receptor is in a red-grey-blue gradient. Orientation: extracellular up, TM3 (left) or TM6 (right) in front.

**Figure 4.**
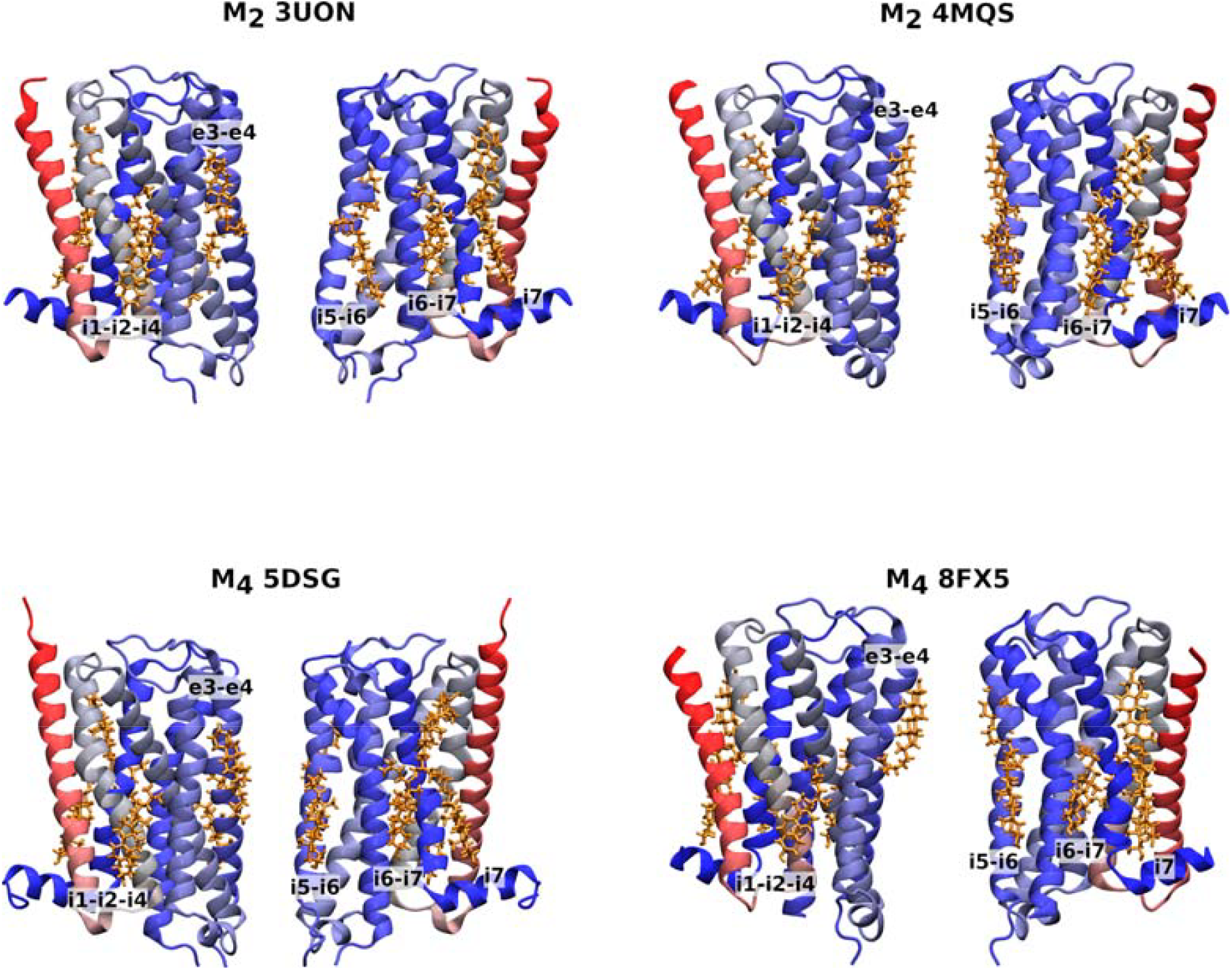
Cholesterol binding to muscarinic receptors. Molecules of cholesterol that are associated with opioid and muscarinic receptors in an inactive or active conformation during cgMD were back-mapped and re-docked to the receptors. Cholesterol is gold. The receptor is in a red-grey-blue gradient. Orientation: extracellular up, TM3 (left) or TM6 (right) in front.

**Figure 5.**
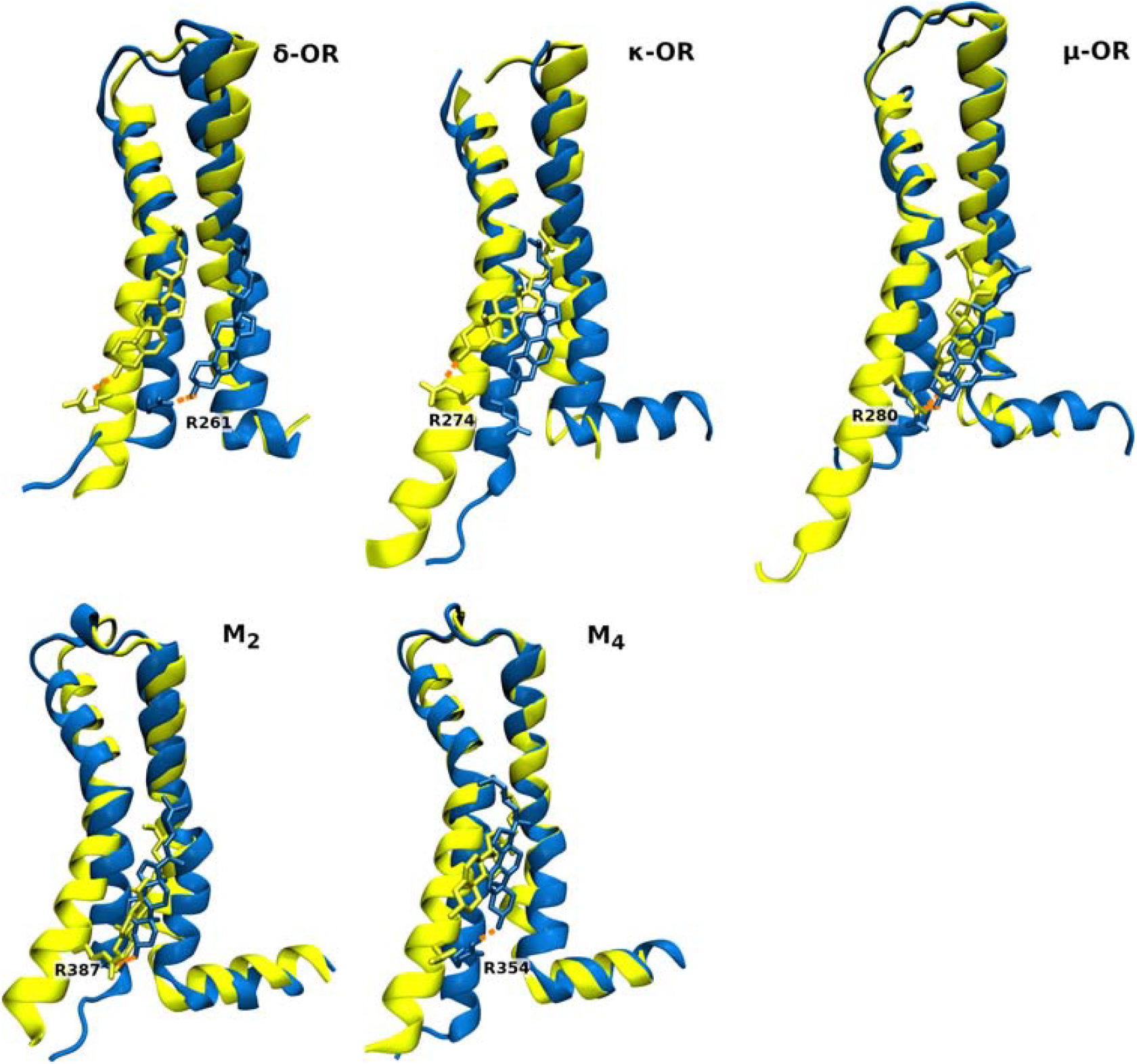
Interaction of cholesterol with R^6.35^ in top docking poses. Interaction of cholesterol with R^6.35^ in top top-scoring docking pose of opioid (left) and muscarinic receptors (right) in an active (yellow) and an inactive conformation (blue). Orange dashed lines indicate hydrogen bonds.

### 3.3 Conventional molecular dynamics

The complexes of receptors with the top-scoring cholesterol molecule in re-docking were subjected to three replicas of conventional MD (cMD). Analysis of cMD trajectories revealed various degrees of stability of bound cholesterol. The quantitative analysis of interactions between receptor and cholesterol molecules is shown in Figure 6. Fractions of time-frames at which cholesterol binds to a given residue were calculated in the Maestro Simulation Interactions Diagram. Residues interacting with cholesterol in at least 20 % of time frames in at least one replica are shown. The principal (the most frequent in the site) receptor residues interacting with cholesterol are listed in Table 2. Differences in cholesterol binding to receptors in inactive versus active conformations were detected.

**Table 1.** List of receptor structures used in molecular modelling. The list of receptor structures includes their activity state, PDB ID code, the method used to determine the structure, structure resolution, number of co-crystallised molecules of cholesterol per receptor molecule and source reference.

| Receptor | Activity | PDB ID | Method | Resolution<br>[Å] | Crystallised<br>cholesterol | Ref. |
| --- | --- | --- | --- | --- | --- | --- |
| A<sub>2A</sub>-adenosine | inactive | 5IU4 | X-ray | 1.72 | 4 | [32] |
| β<sub>2</sub>-adrenergic | inactive | 3D4S | X-ray | 2.80 | 2 | [33] |
| M<sub>2</sub>-muscarinic | inactive | 3UON | X-ray | 3.00 | 0 | [34] |
| M<sub>2</sub>-muscarinic | active | 4MQS | X-ray | 3.50 | 0 | [35] |
| M<sub>4</sub>-muscarinic | inactive | 5DSG | X-ray | 2.60 | 0 | [36] |
| M<sub>4</sub>-muscarinic | active | 8FX5 | Cryo-EM | 2.60 | 0 | [37] |
| δ-opioid | inactive | 4EJ4 | X-ray | 3.4 | 0 | [38] |
| δ-opioid | active | 6PT2 | X-ray | 2.8 | 1 | [39] |
| κ-opioid | inactive | 4DJH | X-ray | 2.90 | 0 | [40] |
| κ-opioid | active | 6B73 | X-ray | 3.10 | 1 | [41] |
| μ-opioid | inactive | 4DKL | X-ray | 2.8 | 1 | [42] |
| μ-opioid | active | 5C1M | X-ray | 2.07 | 1 | [43] |

**Table 2.** Principal residues interacting with cholesterol during conventional MD. Principal residues of interaction between receptor (the first column) and cholesterol molecule in the extracellular leaflet interacting with TM1-TM2-TM7 (e1-e2-e7) bundle or TM4-TM5 (e4-e5) or TM6-TM7 (e6-e7) groove and in the intracellular leaflet interacting with TM1-TM2-TM4 (i1-i2-i4) bundle or TM3-TM4 (i3-i4), TM5-TM6 (i5-i6) or TM6-TM7 (i6-i7) groove or TM7 (i7) helix throughout MD trajectory are listed. Superscript, Ballesteros-Weinstein numbering; ECL3, the third extracellular loop; ICL3, the third intracellular loop.

| Structure | e1-e2-e7 | e4-e5 | e6-e7 | i1-i2-i4 | i3-i4 | i5-i6 | i6-i7 | i7 |
| --- | --- | --- | --- | --- | --- | --- | --- | --- |
| δOR 4EJ4<br>inactive | L295<sup>7.30</sup> | | F280<sup>6.54</sup><br>D293<sup>7.28</sup> | | F137<sup>3.41</sup><br>Y147<sup>3.51</sup> | | R261<sup>6.35</sup><br>V265<sup>6.39</sup><br>F320<sup>7.55</sup> | |
| δOR 6PT2<br>active | S44<sup>1.27</sup> | | F280<sup>6.54</sup><br>D288<sup>6.62</sup> | | K155<sup>3.59</sup> | | R261<sup>6.35</sup><br>G268<sup>6.42</sup> | |
| κOR 4DJH<br>inactive | Y119<sup>2.64</sup><br>S310<sup>7.33</sup><br>Y313<sup>7.36</sup><br>F314<sup>7.37</sup> | | F293<sup>6.54</sup><br>S311<sup>7.34</sup> | | F147<sup>3.41</sup><br>K165<sup>3.59</sup> | | R274<sup>6.35</sup> | |
| κOR 6B73<br>active | S310<sup>7.33</sup><br>F314<sup>7.37</sup> | | F239<sup>6.54</sup><br>S305<sup>7.28</sup><br>S311<sup>7.34</sup> | | | | R274<sup>6.35</sup><br>L325<sup>7.48</sup><br>F332<sup>7.55</sup> | |
| μOR 4DKL<br>inactive | F320<sup>7.37</sup><br>A323<sup>7.40</sup> | | Y299<sup>6.54</sup><br>T311<sup>7.28</sup><br>Q314<sup>ECL3</sup> | | | Y252<sup>5.59</sup> | R280<sup>6.35</sup><br>L331<sup>7.48</sup><br>F338<sup>7.55</sup> | V334<sup>7.51</sup><br>A337<sup>7.54</sup><br>R348<sup>Helix8</sup> |
| μ-OR<br>5C1M<br>active | S317<sup>7.33</sup><br>F320<sup>7.37</sup> | | Y299<sup>6.54</sup><br>Q314<sup>ECL3</sup> | | | I256<sup>5.63</sup><br>K260<sup>I<sub>CL</sub>3</sup> | R280<sup>6.35</sup> | V334<sup>7.51</sup><br>F338<sup>7.55</sup> |
| M<sub>2</sub> 3UON<br>inactive | I26<sup>1.35</sup><br>Y84<sup>2.64</sup><br>W427<sup>7.40</sup> | W155<sup>4.57</sup><br>F180<sup>5.32</sup><br>T190<sup>5.42</sup> |  | Y60<sup>2.41</sup><br>F63<sup>2.44</sup><br>T137<sup>4.39</sup><br>W148<sup>4.50</sup> |  | Y206<sup>5.58</sup><br>E382<sup>6.30</sup> | R387<sup>6.35</sup><br>A391<sup>6.39</sup><br>L394<sup>6.42</sup><br>L442<sup>7.55</sup> | K448<sup>Helix8</sup> |
| M<sub>2</sub> 4MQS<br>active | I26<sup>1.35</sup><br>Y84<sup>2.64</sup><br>W427<sup>7.40</sup> |  |  | K49<sup>1.58</sup><br>Y60<sup>2.41</sup><br>F63<sup>2.44</sup><br>W148<sup>4.50</sup> |  | S210<sup>5.63</sup><br>T386<sup>6.34</sup><br>R387<sup>6.35</sup> | A391<sup>6.39</sup><br>A438<sup>7.51</sup><br>K448<sup>Helix8</sup> | L442<sup>7.55</sup><br>K448<sup>Helix8</sup><br>K452<sup>Helix8</sup> |
| M<sub>4</sub> 5DSG<br>inactive | E31<sup>1.31</sup><br>Y92<sup>2.64</sup><br>S290<sup>7.36</sup> | W164<sup>4.57</sup><br>T199<sup>5.42</sup> |  | F72<sup>2.44</sup><br>W157<sup>4.50</sup> |  | S219<sup>5.63</sup> | R354<sup>6.35</sup> | A405<sup>7.51</sup><br>A408<sup>7.54</sup><br>K415<sup>Helix8</sup> |
| M<sub>4</sub> 8FX5<br>active | F34<sup>1.35</sup><br>W440<sup>7.40</sup> |  |  | F72<sup>2.44</sup><br>W157<sup>4.50</sup> |  | S219<sup>5.63</sup> | R400<sup>7.46</sup> | K415<sup>Helix8</sup><br>R419<sup>Helix8</sup> |

**Figure 6.**
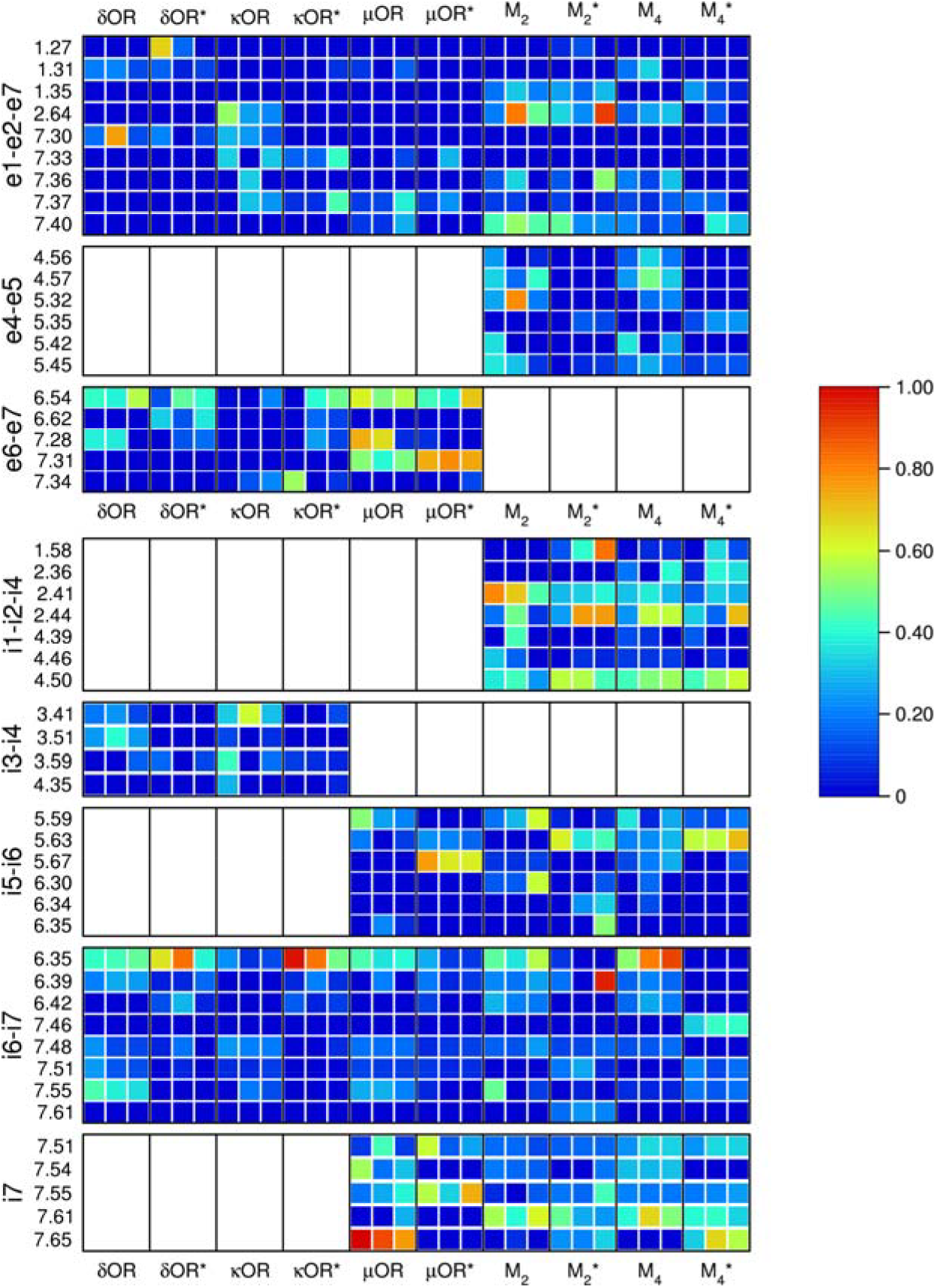
Interactions between cholesterol and receptor. Histograms of interactions between receptors indicated on the x-axis and cholesterol molecules in the extracellular leaflet interacting with e1-e2-e7 bundle, e4-e5 groove or e6-e7 groove and in the intracellular leaflet interacting with i1-i2-i4 bundle, i3-i4 groove, i5-i6 groove, i6-i7 groove or i7 helix indicated on the y-axis were calculated from MD trajectories using Ligand Interaction Diagram in Maestro. Only residues interacting with cholesterol at more than 20% of time frames in at least one MD replica are displayed. Each square represents one of 3 independent MD replicas. Values over 1.0 are possible as some protein residues may make multiple contacts with cholesterol within one timeframe. However, no values greater than 1 were obtained.

#### 3.3.1 Opioid receptors

Four unique cholesterol-binding sites were identified on the δ-opioid receptor (Table 2, Figure 3). Binding to e6-e7 and i3-i4 sites was similar for receptors at inactive and active conformations, respectively. In the e6-e7 site, the cholesterol binding was mediated by hydrophobic interaction with F280^6.54^ (superscript, Ballesteros-Weinstein numbering[27]) (Figure 6, Table 2). Cholesterol binding to the i3-i4 site was weak at both conformations. In an inactive conformation (4EJ4), binding was mediated by hydrophobic interactions with F137^3.41^. In an active conformation (6PT2), binding was mediated by hydrogen bonding to K155^3.59^. While cholesterol binding to the e1-e2-e7 site at an inactive conformation was primarily mediated by hydrophobic interaction with L295, at an active conformation, it was mediated by hydrogen bonding to S44 ^1.27^. Cholesterol binding to the i6-i7 site of the δ-opioid receptor was via a hydrogen bond to R261^6.35^ that was more frequent in an active conformation than in an inactive conformation. In an active conformation, cholesterol was shifted towards TM7, interacting with F320^7.55^ (Figure 5).

Similar to the δ-opioid receptor, four unique cholesterol-binding sites were identified on the κ-opioid receptor (Table 2, Figure 3). In contrast to the δ-opioid receptor, however, cholesterol binding to the extracellular e1-e2-e7 site was more frequent at an inactive conformation (4DJH) than at an active conformation (6B73) (Figure 6). In an inactive conformation, it was mediated by frequent hydrogen bonding with Y119^2.64^ and S310^7.33^, and in an active conformation by less-frequent hydrogen bonding with S310^7.33^ only. Cholesterol binding to the e6-e7 site was mediated by hydrogen bonding to S311^7.34^ and hydrophobic interactions with F293^6.54^ that were more frequent in an active than inactive conformation. Cholesterol binding to the i3-i4 site of the receptor in an inactive conformation was mediated by F147^3.41^ and K165^3.59^. At the receptor in an active conformation, cholesterol binding was weak and cholesterol dissociated from the binding site. In the i6 site, the cholesterol binding was primarily mediated via a hydrogen bond with R274^6.35^, which was more frequent at an active conformation than an inactive one (Figure 5).

Five unique cholesterol-binding sites were identified on the μ-opioid receptor (e1-e2-e7, e6-e7, i3-i4, i6-i7, i7), three being common with δ- and κ-opioid receptors (Table 2, Figure 3). The strongest cholesterol binding was identified at the i7 site (Figure 6, Table 2). In an inactive conformation (4DKL), cholesterol binding to the i7 site was mediated by R348 in Helix8. In an active conformation (5C1M), cholesterol binding to the i7 site was mediated by V334^7.51^ and F338^7.55^. The cholesterol binding to the i7 site was more frequent in an inactive conformation than in an active one. Also, binding to the e6-e7 site was relatively strong. It was mediated by hydrophobic interaction with Y299^6.54^ and hydrogen bonding to Q314 in the third extracellular loop. At inactive conformation, hydrogen bonding to Q314 was less frequent in expense for hydrogen bonding to T311^7.28^. Cholesterol binding to the i5-i6 site was mediated by hydrogen bonding to K260 in the third intracellular loop of an active conformation or less frequent binding to Y252^5.59^ in the TM5 helix of an inactive conformation. Like at other tested receptors, cholesterol binding to the i6-i7 site was mediated via a hydrogen bond with R280^6.35^, interacting more frequently with it at an inactive conformation (Figure 5). In an inactive conformation, cholesterol binding was further stabilised by hydrophobic interactions with TM7 via L331^7.48^ and F338^7.55^ (Figure 6).

#### 3.3.2 Muscarinic receptors

Six unique cholesterol-binding sites were identified on muscarinic receptors (Table 2, Figure 4). Interaction patterns of M_2_ and M_4_ receptors were similar. The most profound differences between inactive and active conformations were observed for cholesterol binding to the e4-e5 and i6-i7 sites (Figure 6). In an inactive conformation of the M_2_-muscarinic receptor (3UON), cholesterol at the e4-e5 site primarily interacted with W155^4.57^ and T190^5.42^. Identically, in an inactive conformation of the M_4_-muscarinic receptor (5DSG), cholesterol primarily interacted with W164^4.57^ and T199^5.42^. In active conformations (M_2_ 4MQS; M_4_ 8FX5), these principal interactions were missing as cholesterol binding was unstable. Like opioid receptors, cholesterol binding to the i6-i7 site of the M_2_ and M_4_ receptors in inactive conformations was mainly mediated by hydrogen bonding to R387^6.35^ and R354^6.35^, respectively. In active conformations, cholesterol moved towards Helix8 and bound to K448 of M_2_ or R400 of M_4_ receptor. Cholesterol binding to the e1-e2-e7 site of the M_2_ receptor in an inactive conformation was mainly mediated by Y84^2.64^ and W427^7.40^. In an active conformation, these interactions persisted, but bonding to W427^7.40^ was about 5 times less frequent. Cholesterol binding to the e1-e2-e7 site of the M_4_ receptor was, in principle, the same but much weaker and at an inactive conformation of the receptor unstable. Cholesterol binding to the i1-i2-i4 site of the M_2_ and M_4_ receptors was strong. At the M_2_-receptor, cholesterol binding was mainly mediated by Y60^2.41^, F63^2.44^, T237^4.39^ and W148^4.50^ at an inactive conformation and K49^1.58^, Y60^2.41^, F63^2.44^ and W148^4.50^ at an active conformation. At the M_4_-receptor, cholesterol binding to the i1-i2-i4 site was mainly mediated by F63^2.44^ and W148^4.50^ at both conformations. Cholesterol binding to the i5-i6 site of the M_2_ receptor was mainly mediated by Y206^5.58^ and E382^6.30^ at an inactive conformation and by S210 ^5.63^, T386^6.34^ at an active conformation. In an active conformation, cholesterol moves towards R387^6.35^ and forms hydrogen bonds with it (Figure 5). Cholesterol binding to the i5-i6 site of the M_4_ receptor was mainly mediated by S219^5.63^ at both conformations (Figure 6). Cholesterol binding to the i7 site was mediated by hydrogen bonding to lysine (M_2_ K448, M_4_ K415)in Helix 8 at both active and inactive conformations. At active conformations, it moves further from the TM7 helix and forms a hydrogen bond with K452 (M_2_) or R419 (M_4_).

#### 3.3.3 Common features of cholesterol binding

Taken together, the sites e1-e2-e7 and i6-i7 were detected in all structures (Table 2, Figure 6). Cholesterol binding to the e1-e2-e7 bundle varied among receptor families and subtypes. Cholesterol binding to the e1-e2-e7 bundle was weak at μ-opioid and M_4_-muscarinic receptors. Cholesterol binding to the e1-e2-e7 bundle was mediated mainly by Y^2.64^ at M_2_-muscarinic and κ-opioid receptors, by L^7.30^ at δ-opioid receptor in an inactive conformation and S^1.27^ at δ-opioid receptor in an active conformation. The principal interaction residue of cholesterol binding to the i6-i7 groove was R^6.35^ (Figure 5). Cholesterol binding to R^6.35^ was more frequent at inactive than active conformations of M_2_, M_4_ and μ receptors (Figure 6). In contrast, cholesterol binding to R^6.35^ was more frequent to active conformations than to inactive conformations of κ and δ-opioid receptors. Cholesterol binding to the i6-i7 site diminishes the dynamic movement of TM6 as evidenced by lower RMSF (Figure 7). RMSF of TM6 with cholesterol bound to the i6-i7 site was smaller in inactive than active conformation of M_2_ and M_4_ receptors by 26 and 15 %, respectively, but the same for both muscarinic receptors in active conformations. In contrast, the RMSF of TM6 with cholesterol bound to the i6-i7 site was smaller at the active than inactive conformation of κ and μ receptors by 28 and 25 %, respectively, but the same for these receptors in inactive conformations. Cholesterol binding to the i6-i7 site of δ receptor did not affect the dynamics of TM6 in an inactive or active conformation.

**Figure 7.**
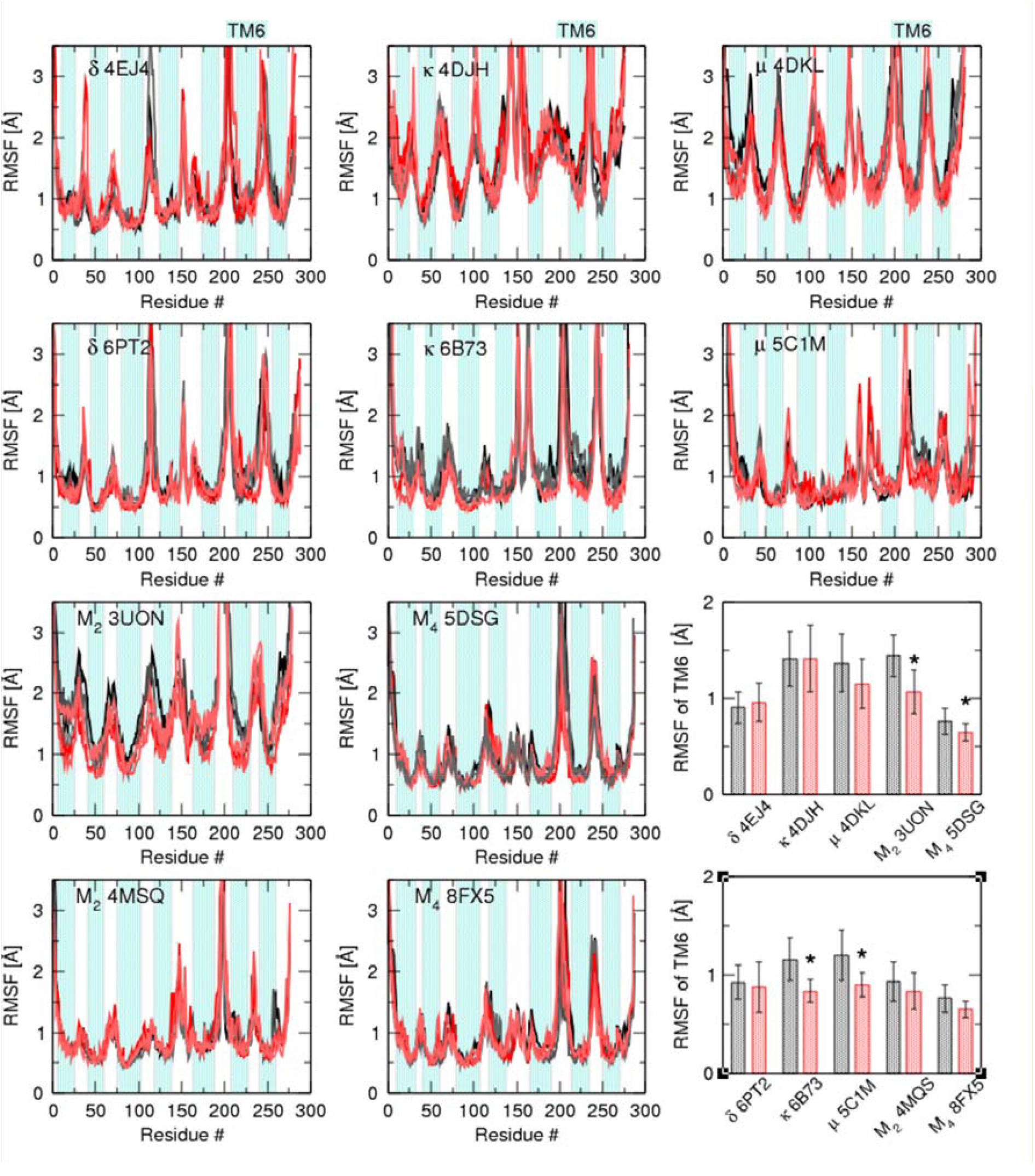
Receptor RMSF during MD simulation. Root mean square fluctuations (RMSF) in Å of individual residues of receptors indicated in legends in the absence (shades of grey) or presence (shades of red) of cholesterol bound to the i6-i7 site are plotted against residue number. Traces are from 3 independent MD replicas. Turquoise backgrounds indicate individual transmembrane helices. Lower right, bar plots of RMSF of TM6 are means ± SD. *, different from apo structure (P<0.05 according to t-test with Bonferroni correction for multiple comparisons.

### 3.4 Functional responses – membrane-cholesterol depletion

To assess the effects of membrane cholesterol on the functional response of muscarinic and opioid receptors, membrane cholesterol was depleted in CHO cells expressing individual receptor subtypes by treatment with methyl-β-cyclodextrin (MβCD). The treatment lowered the concentration of membrane cholesterol from 225 ± 22 to 85 ± 13 nmol/mg of protein. The cholesterol removal by this procedure was shown to be reversible[28, 56]. Such a drastic decrease in cholesterol concentration is needed to remove cholesterol from receptor binding, as cholesterol may exert high affinity, up to 1 nM[57]. Cholesterol depletion did not change receptor binding capacity. No novel sites were exposed or lost by MβCD treatment. To avoid the effects of cholesterol removal on downstream signalling membrane proteins, functional response was measured close to the receptor as [^35^S]GTPγS binding to membranes prepared from CHO cells. The M_2_ and M_4_-muscarinic receptors were activated by the agonist carbachol, δ- and κ-opioid receptors by Leu-enkephalin and μ-opioid receptors by endomorphin I. After subtraction of non-specific binding, [^35^S]GTPγS binding was normalised to total GTPγS binding sites. The effects of cholesterol depletion varied among receptors. Cholesterol depletion changed basal GTPγS binding (% of total GTPγS binding sites in the absence of an agonist) to membranes, increasing it at M_2_- and M_4_-muscarinic receptors by 18% (from 2.46 to 2.9%) (Figure 8, red and turquoise) and decreasing it at δ- and μ-opioid receptors by 13% (from 1.92 to 1.68%) and 44% (from 2.08 to 1.18%), respectively (Figure 8, yellow and blue, Table 3). Basal GTPγS binding to membranes expressing κ-opioid receptors remained unchanged. Membrane-cholesterol depletion increased maximum GTPγS binding at both muscarinic receptors by about 20 % and decreased maximum GTPγS binding at all opioid receptors by 20 (μ-opioid) to 50 % (κ-opioid). Membrane-cholesterol depletion decreased the half-efficient concentration EC_50_ at both muscarinic 1.5-fold at M_2_ (5.9 μM vs. 9.1 μM) and 2.7-fold at M_4_ (1.9 μM vs. 5.2 μM) and μ-opioid receptors 1.4-fold (12.8 nM vs. 17.8 nM) and increased it at κ-opioid receptor 1.8-fold (27 μM vs 15μM). EC_50_ was not affected at δ-opioid receptors (Table 3). Agonists used did not affect GTPγS binding to membranes from non-transfected cells, demonstrating that GTPγS binding is specific to activation of transfected receptors.

**Table 3.** Parameters of functional response to agonists at native and methyl-β-cyclodextrin-treated (MβCD) membranes. EC_50_ values are expressed as negative decadic logarithms. E_MAX_ and basal values are expressed as % of occupancy of G-protein binding sites. Data are means ± SD from 3 independent experiments performed in quadruplicates.

|  | pEC<sub>50</sub> | E<sub>MAX</sub><br>[% occupancy] | Basal<br>[% occupancy] |
| --- | --- | --- | --- |
| δ-OR | 7.87 ± 0.08 | 10.65 ± 0.38 | 1.92 ± 0.07 |
| δ-OR MβCD | 7.86 ± 0.02 | 7.35 ± 0.07* | 1.68 ± 0.06* |
| κ-OR | 4.82 ± 0.06 | 10.10 ± 0.29 | 2.50 ± 0.08 |
| κ-OR MβCD | 4.57 ± 0.05* | 5.27 ± 0.11* | 2.40 ± 0.08 |
| μ-OR | 7.75 ± 0.05 | 11.55 ± 0.28 | 2.09 ± 0.07 |
| μ-OR MβCD | 7.89 ± 0.07* | 9.38 ± 0.31* | 1.18 ± 0.05* |
| M<sub>2</sub> | 5.04 ± 0.04 | 12.14 ± 0.26 | 2.46 ± 0.08 |
| M<sub>2</sub> MβCD | 5.23 ± 0.04* | 14.97 ± 0.25* | 2.90 ± 0.09* |
| M<sub>4</sub> | 5.28 ± 0.07 | 11.13 ± 0.35 | 2.46 ± 0.08 |
| M<sub>4</sub> MβCD | 5.72 ± 0.02* | 13.19 ± 0.10* | 2.90 ± 0.09* |

\*, different from untreated ( $P < 0.05$ , paired t-test)

**Figure 8.**
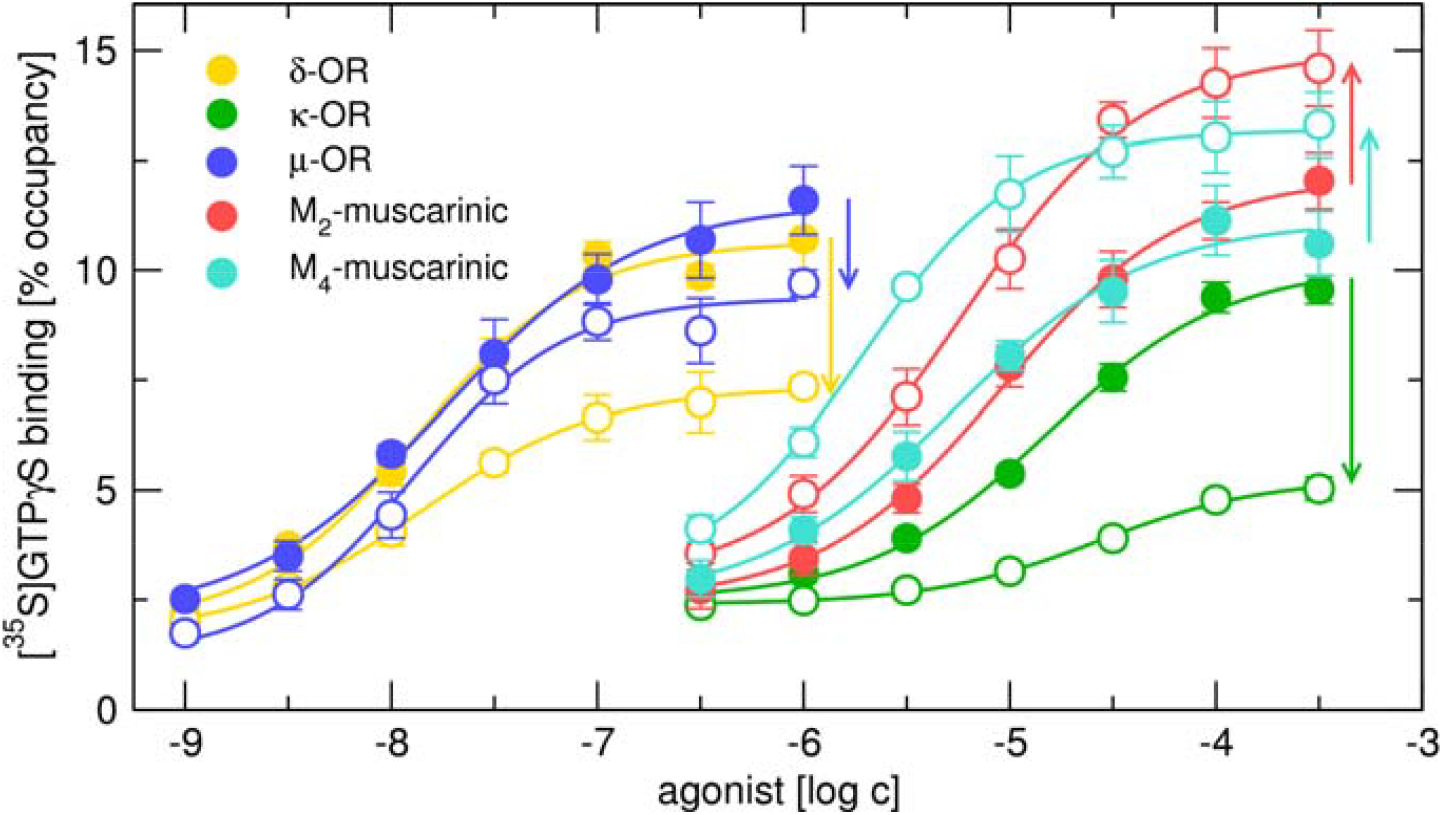
Effects of membrane-cholesterol depletion on functional response to agonists. Effects of membrane-cholesterol depletion (open symbols) on functional responses of δ- (yellow), κ-opioid (green) receptors to Leu-enkephalin, μ-opioid receptors (blue) to endomorphin I and M_2_ (red) and M muscarinic (turquoise) receptors to carbachol. Ordinate, [^35^S]GTPγS binding is expressed as % of occupancy of G-protein binding sites. Abscissa, the concentration of agonist is expressed as a decadic logarithm of molar concentration. Data are means ± SD from 3 independent experiments performed in quadruplicates.

### 3.5 Functional responses – mutations

Molecular modelling (Figure 6) suggests that cholesterol binds to the i6-i7 groove at all investigated receptors and R plays a key role in cholesterol binding at the majority of them. Therefore, R was mutated to either alanine (neutral mutation) or glutamate (reverse charge mutation) found in some receptors (Figure 1) with the intention of abolishing cholesterol binding to R. The functional response of M_2_- and M_4_-muscarinic and δ-, κ- and μ-opioid receptors was measured as stimulation of [ S]GTPγS binding to membranes prepared from CHO cells by the agonists carbachol (muscarinic), Leu-enkephalin (δ- and κ-opioid) or endomorphin I (μ-opioid receptors). The mutations of R affected functional responses similarly to cholesterol depletion. Both mutations increased basal binding (in the absence of an agonist) of GTPγS binding to M_2_- and M_4_-muscarinic receptors (Figure 9, red and turquoise) and decreased basal GTPγS binding at δ- and μ-opioid receptors (Figure 9, yellow and blue). None of the mutations affected basal GTPγS binding to membranes expressing κ-opioid receptors (Figure 9, green). Both mutations increased maximum GTPγS binding at both muscarinic receptors and decreased maximum GTPγS binding at the κ-opioid receptor. Mutation of R to glutamine decreased E_MAX_ at μ-opioid receptor. Mutations of R to alanine did not affect E_MAX_ at this receptor. Both mutations decreased EC_50_ values at both muscarinic and κ-opioid receptors. EC_50_ was not affected at δ- and μ-opioid receptors by these mutations (Table 4).

**Table 4.**
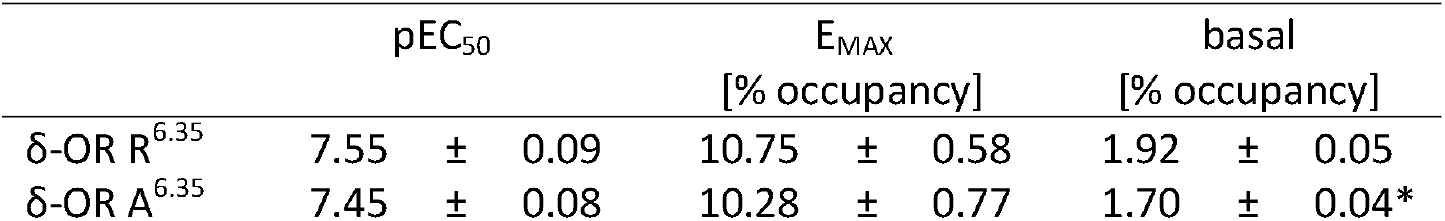
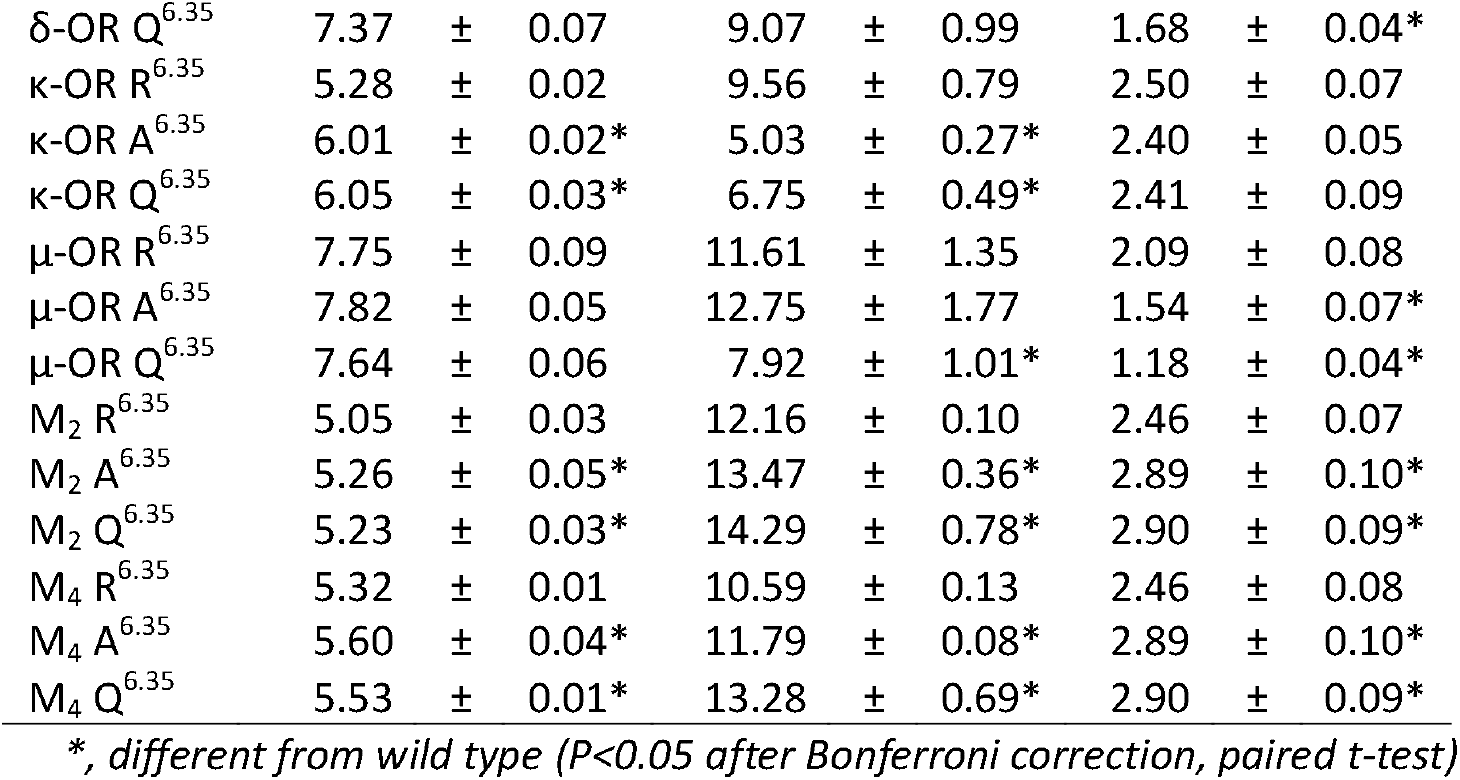
Parameters of functional response to agonists at wild type (R^6.35^) and mutated receptors. EC_50_ values are expressed as negative decadic logarithms. E_MAX_ and basal values are expressed as % of occupancy of G-protein binding sites. Data are means ± SD from 3 independent experiments

| | $pEC_{50}$ | $E_{\text{MAX}}$<br>[% occupancy] | basal<br>[% occupancy] |
| --- | --- | --- | --- |
| $\delta\text{-OR } R^{6.35}$ | 7.55 ± 0.09 | 10.75 ± 0.58 | 1.92 ± 0.05 |
| $\delta\text{-OR } A^{6.35}$ | 7.45 ± 0.08 | 10.28 ± 0.77 | 1.70 ± 0.04* |

|  |  |  |  |  |  |  |  |  |  |
| --- | --- | --- | --- | --- | --- | --- | --- | --- | --- |
| δ-OR Q<sup>6.35</sup> | 7.37 | ± | 0.07 | 9.07 | ± | 0.99 | 1.68 | ± | 0.04* |
| κ-OR R<sup>6.35</sup> | 5.28 | ± | 0.02 | 9.56 | ± | 0.79 | 2.50 | ± | 0.07 |
| κ-OR A<sup>6.35</sup> | 6.01 | ± | 0.02* | 5.03 | ± | 0.27* | 2.40 | ± | 0.05 |
| κ-OR Q<sup>6.35</sup> | 6.05 | ± | 0.03* | 6.75 | ± | 0.49* | 2.41 | ± | 0.09 |
| μ-OR R<sup>6.35</sup> | 7.75 | ± | 0.09 | 11.61 | ± | 1.35 | 2.09 | ± | 0.08 |
| μ-OR A<sup>6.35</sup> | 7.82 | ± | 0.05 | 12.75 | ± | 1.77 | 1.54 | ± | 0.07* |
| μ-OR Q<sup>6.35</sup> | 7.64 | ± | 0.06 | 7.92 | ± | 1.01* | 1.18 | ± | 0.04* |
| M<sub>2</sub> R<sup>6.35</sup> | 5.05 | ± | 0.03 | 12.16 | ± | 0.10 | 2.46 | ± | 0.07 |
| M<sub>2</sub> A<sup>6.35</sup> | 5.26 | ± | 0.05* | 13.47 | ± | 0.36* | 2.89 | ± | 0.10* |
| M<sub>2</sub> Q<sup>6.35</sup> | 5.23 | ± | 0.03* | 14.29 | ± | 0.78* | 2.90 | ± | 0.09* |
| M<sub>4</sub> R<sup>6.35</sup> | 5.32 | ± | 0.01 | 10.59 | ± | 0.13 | 2.46 | ± | 0.08 |
| M<sub>4</sub> A<sup>6.35</sup> | 5.60 | ± | 0.04* | 11.79 | ± | 0.08* | 2.89 | ± | 0.10* |
| M<sub>4</sub> Q<sup>6.35</sup> | 5.53 | ± | 0.01* | 13.28 | ± | 0.69* | 2.90 | ± | 0.09* |

\*, different from wild type ( $P < 0.05$ after Bonferroni correction, paired $t$ -test)

**Figure 9.**
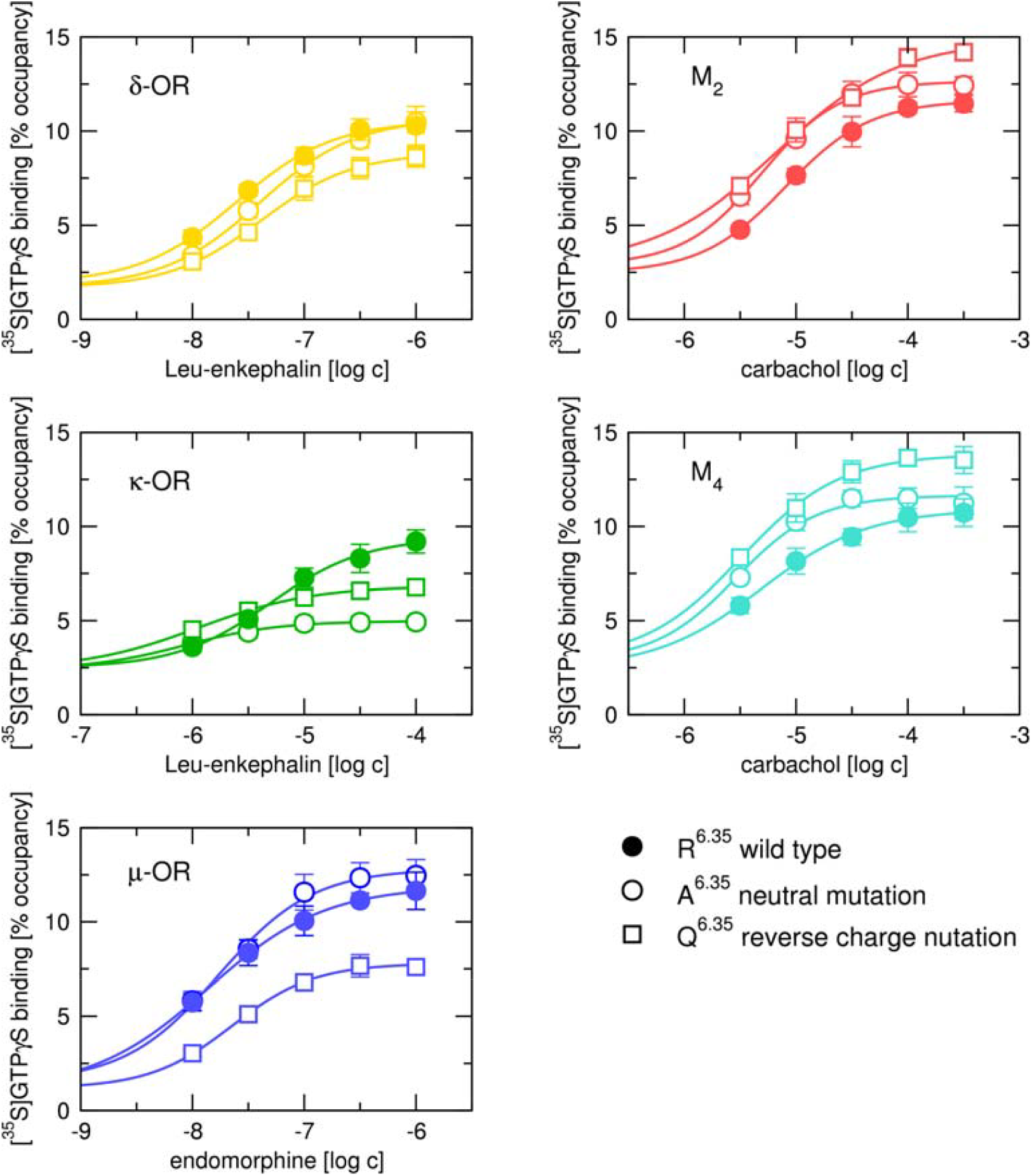
Effects of mutation of R^6.35^ on functional response to agonists. Effects of mutations of wild type R^6.35^ (full symbols) to alanine (neutral mutation, open circles) or glutamine (reverse charge mutation, open squares) on functional responses of δ- (yellow), κ-opioid (green) receptors to Leu-enkephalin, μ-opioid receptors (blue) to endomorphin I and M_2_ (red) and M_4_muscarinic (turquoise) receptors to carbachol. Ordinate, [^35^S]GTPγS binding is expressed as % of occupancy of G-protein binding sites. Abscissa, the concentration of agonist is expressed as a decadic logarithm of molar concentration. Data are means ± SD from 3 independent experiments performed in quadruplicates.

Overall, the effects of mutations on basal GTPγS binding were the same as the effects of cholesterol depletion. Also, the effects of mutations on E_MAX_ and EC_50_ at muscarinic receptors were the same as the effects of cholesterol depletion. In contrast, cholesterol depletion strongly decreased the E_MAX_ of functional response of mutated M_2_- and M_4_-muscarinic receptors, having the opposite effect in comparison to wild-type receptors (Figure 10 and Table 5), indicating an additional mechanism by which cholesterol affects functional properties of muscarinic receptors.

**Table 5.** Parameters of functional response to agonists at mutated receptors. EC_50_ values are expressed as negative decadic logarithms. E_MAX_ and basal values are expressed as % of occupancy of G-protein binding sites. Data are means ± SD from 3 independent experiments

|  | pEC<sub>50</sub> |  |  | E<sub>MAX</sub><br>[% occupancy] |  |  | basal<br>[% occupancy] |  |  |
| --- | --- | --- | --- | --- | --- | --- | --- | --- | --- |
| M<sub>2</sub> A<sup>6.35</sup> | 5.26 | ± | 0.02 | 13.55 | ± | 0.29 | 2.90 | ± | 0.04 |
| M<sub>2</sub> A<sup>6.35</sup> MβCD | 6.17 | ± | 0.03* | 4.46 | ± | 0.20* | 2.88 | ± | 0.04 |
| M<sub>2</sub> Q<sup>6.35</sup> | 5.26 | ± | 0.02 | 14.07 | ± | 1.01 | 2.94 | ± | 0.01 |
| M<sub>2</sub> Q<sup>6.35</sup> MβCD | 5.73 | ± | 0.01* | 4.50 | ± | 0.22* | 2.89 | ± | 0.03 |
| M<sub>4</sub> A<sup>6.35</sup> | 5.64 | ± | 0.02 | 11.71 | ± | 0.93 | 2.89 | ± | 0.04 |
| M<sub>4</sub> A<sup>6.35</sup> MβCD | 6.50 | ± | 0.02* | 5.11 | ± | 0.01* | 2.71 | ± | 0.02* |
| M<sub>4</sub> Q<sup>6.35</sup> | 5.50 | ± | 0.02 | 13.40 | ± | 0.60 | 2.92 | ± | 0.02 |
| M<sub>4</sub> Q<sup>6.35</sup> MβCD | 5.89 | ± | 0.02* | 5.22 | ± | 0.10* | 2.82 | ± | 0.02* |

\*, different from untreated ( $P < 0.05$ , paired t-test)

**Figure 10.**
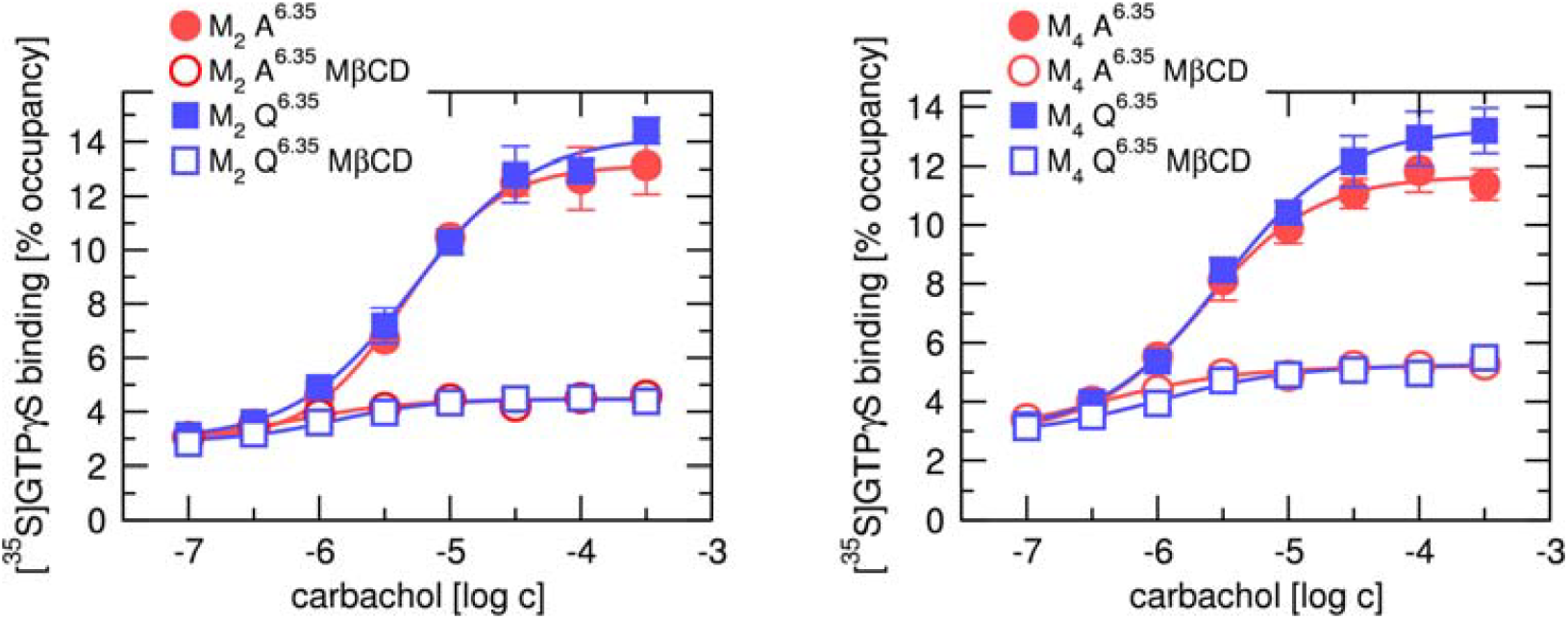
Effects of cholesterol depletion on the functional response of mutated muscarinic receptors. Effects of membrane-cholesterol depletion (open symbols) on functional responses of mutated (red circles - R^6.35^ to alanine; blue squares – R^6.35^ to glutamine) M_2_ (left) and M_4_ muscarinic (right) receptors to carbachol. Ordinate, [^35^S]GTPγS binding is expressed as % of occupancy of G-protein binding sites. Abscissa, the concentration of agonist is expressed as a decadic logarithm of molar concentration. Data are means ± SD from 3 independent experiments performed in quadruplicates.

### 3.6 Receptor expression level

As parameters of functional responses (EC_50_ and E_MAX_) are dependent on the receptor expression level, the maximum binding capacity (B_MAX_) and equilibrium dissociation constant (K_D_) of radioligand were determined in saturation binding experiments (Table 6). In general, mutations of R did not bring substantial changes in receptor B_MAX_ or radioligand K_D_. Only mutations R261Q of δ-OR and R280A and R280Q of μ-OR marginally increased receptors B_MAX_ (<7 %).

**Table 6.** Expression of muscarinic and opioid receptors. Maximum binding capacity B_MAX_, expressed as fmol of binding sites per μg of membrane proteins, equilibrium dissociation constant KD, expressed in nM, of [^3^H]NMS (muscarinic), [^3^H]DAMGO (μ-OR), [^3^H]delthropin (δ-OR) and [^3^H]diprenorphine (κ-OR), respectively, of wild type (R^6.35^) and mutant receptors were obtained by fitting Eq. 2 to saturation binding data. Data are means ± SD from 3 independent experiments performed in quadruplicates.

| | $B_{MAX}$ [fmol/μg] | | | $K_D$ [nM] | | |
| --- | --- | --- | --- | --- | --- | --- |
| δ-OR R<sup>6.35</sup> | 1.42 | ± | 0.02 | 157 | ± | 5 |
| δ-OR A<sup>6.35</sup> | 1.49 | ± | 0.05 | 163 | ± | 2 |
| δ-OR Q<sup>6.35</sup> | 1.51 | ± | 0.01* | 173 | ± | 29 |
| κ-OR R<sup>6.35</sup> | 1.76 | ± | 0.09 | 1630 | ± | 230 |
| κ-OR A<sup>6.35</sup> | 1.74 | ± | 0.02 | 1440 | ± | 130 |
| κ-OR Q<sup>6.35</sup> | 1.77 | ± | 0.09 | 1240 | ± | 100 |
| μ-OR R<sup>6.35</sup> | 1.49 | ± | 0.01 | 175 | ± | 18 |
| μ-OR A<sup>6.35</sup> | 1.59 | ± | 0.01* | 192 | ± | 4 |
| μ-OR Q<sup>6.35</sup> | 1.57 | ± | 0.06* | 194 | ± | 8 |
| M<sub>2</sub> R<sup>6.35</sup> | 7.50 | ± | 0.02 | 403 | ± | 6 |
| M<sub>2</sub> A<sup>6.35</sup> | 7.59 | ± | 0.12 | 397 | ± | 6 |
| M<sub>2</sub> Q<sup>6.35</sup> | 7.58 | ± | 0.19 | 402 | ± | 3 |
| M<sub>4</sub> R<sup>6.35</sup> | 8.21 | ± | 0.15 | 233 | ± | 3 |
| M<sub>4</sub> A<sup>6.35</sup> | 8.26 | ± | 0.05 | 235 | ± | 5 |
| M<sub>4</sub> Q<sup>6.35</sup> | 8.35 | ± | 0.11 | 237 | ± | 6 |

\*, different from wild-type ( $P < 0.01$ , Bonferroni multiple comparison)

## 4 Discussion

Membrane cholesterol allosterically modulates ligand binding to and activation of GPCRs and affects the pharmacology of GPCRs[2]. Three cholesterol-binding motifs were postulated[15, 16, 22]. However, the majority of cholesterol binding in cryo-EM and X-ray structures of GPCRs occurs in the sites not conforming to these binding motifs[26]. While traditional motif-based predictions may not fully capture the complexity of cholesterol-GPCR interactions, site-specific interactions play a crucial role in modulating GPCR conformations and function. This underscores the need for detailed, residue-level analyses to better understand these interactions. In the present study, we show cholesterol binding to such a non-canonical site at TM6 of muscarinic and opioid receptors. The binding to this site is primarily mediated by hydrogen bonding to R (Figure 2 and Figure 6).

We docked cholesterol to GPCRs by simulation of molecular dynamics of its association with receptors[29]. The procedure successfully identified all cholesterol-binding sites observed in the X-ray structures of the β_2_-adrenergic receptor (3D4S)[22] and A_2A_-adenosine receptor (5IU4)[58], verifying a successful implementation of the procedure. Eight unique cholesterol-binding “cavities” on muscarinic and opioid receptors were identified (Figure 2, Table 2). Two of them were common for muscarinic and opioid receptors. At opioid receptors, observed cholesterol binding in the extracellular leaflet to TM6-TM7 groove (e6-e7 site) corresponds to cholesterol co-crystallised in structures 4DKL, 5C1M, 6PT2 and 6B73, further validating the procedure[59–62]. Cholesterol did not appear in the X-ray or cryo-EM structures of muscarinic receptors. However, the observed binding of cholesterol in the intracellular leaflet to the TM1-TM2-TM4 bundle (i1-i2-i4 site) corresponds to cholesterol hemisuccinate appearing in X-ray (5CXV) and cryo-EM (6OIJ) structures of the M_1_-muscarinic receptor[63, 64]. The i1-i2-i4 site corresponds to the CCM motif[33], however, cholesterol hydrogen bonding to K^4.39^, essential to the CCM motif, was sparse (Figure 6). The binding to the i1-i2-i4 site was mediated mainly by W^4.50^ and Y^2.41^, which are part of the CCM motif and F^2.44^, which is not a part of the CCM motif.

We further analysed the cholesterol binding to the putative sites by simulation of cMD of top-scoring poses (Figure 6). The binding of cholesterol to receptors in active conformations diverged from its binding to receptors in inactive conformations, indicating a modulatory role on receptor activation. Based on changes in receptor dynamics (Figure 7), we assume that preferential binding of cholesterol to one of the receptor conformations leads to stabilisation of the receptor in this conformation and thus influences its activation and, consequently, the functional response. Specifically, preferential binding to the inactive conformation leads to attenuation of the functional response and possibly a decrease in the basal response. On the other hand, preferential binding to the active conformation leads to enhancement of the functional response and possibly an increase in the basal response. We observed major differences in binding to active versus inactive conformation for the i6-i7 site (Figure 5).

Depletion of membrane cholesterol differentially modulated activation of muscarinic and opioid receptors (Figure 8). Depletion of membrane cholesterol augmented activation of both muscarinic receptors to a similar extent and diminished to various extents (20 to 50%) activation of all opioid receptors. These findings agree with previous observations[65]. We assume that the removal of membrane cholesterol diminishes its binding to the receptor, which leads to attenuation of modulatory effects. Thus, the increase in the functional response of muscarinic receptors after the removal of membrane cholesterol means that cholesterol has an overall negative modulatory effect. That is, it binds preferentially to the inactive conformation of the receptor (Figure 6) and stabilises it in an inactive conformation (Figure 7). On the other hand, the decrease in the functional response of opioid receptors means that cholesterol has a positive modulatory effect. That is, it binds preferentially to the active conformation of the receptor (Figure 6) and stabilises it in an active conformation (Figure 7).

Molecular modelling suggested that cholesterol hydrogen bonding to R^6.35^ substantially contributes to its binding to the receptor (Figure 6). Therefore, we designed mutations of R^6.35^ intended to abolish cholesterol hydrogen bonding. Effects of mutations of R^6.35^ to glutamine on functional response to agonists (Figure 9) were similar to the effects of cholesterol depletion (Figure 8), suggesting these mutations indeed impair cholesterol binding to R^6.35^. Thus i6-i7 groove appears to be a “non-canonical” (motif-less) cholesterol binding site through which cholesterol differentially modulates the functional response of studied receptors (muscarinic receptors negatively and opioid receptors positively). The groove is caused by the P^6.50^ kink. Thus, the i6-i7 site may be considered a “tilted” cholesterol-binding motif[21]. The i6-i7 site is located between R^6.35^ and CWXP transmission switch[66]. The R^6.35^ is located opposite to E^6.30^, forming an ionic lock with R^3.50^ of the DRY ionic lock switch. Outward movement of TM6 from the transmembrane helix bundle is a fundamental element of GPCR activation (Figure 11). The binding of cholesterol to R restricts this movement (Figure 7). Therefore, it is reasonable to assume that cholesterol binding to the i6-i7 site stabilises the receptor in a given conformation. The TM6 outward movement in the δ receptor is smaller than in the M_2_ receptor (Figure 11). This may explain the differences observed in the modulation of muscarinic and opioid receptors by cholesterol.

**Figure 11.**
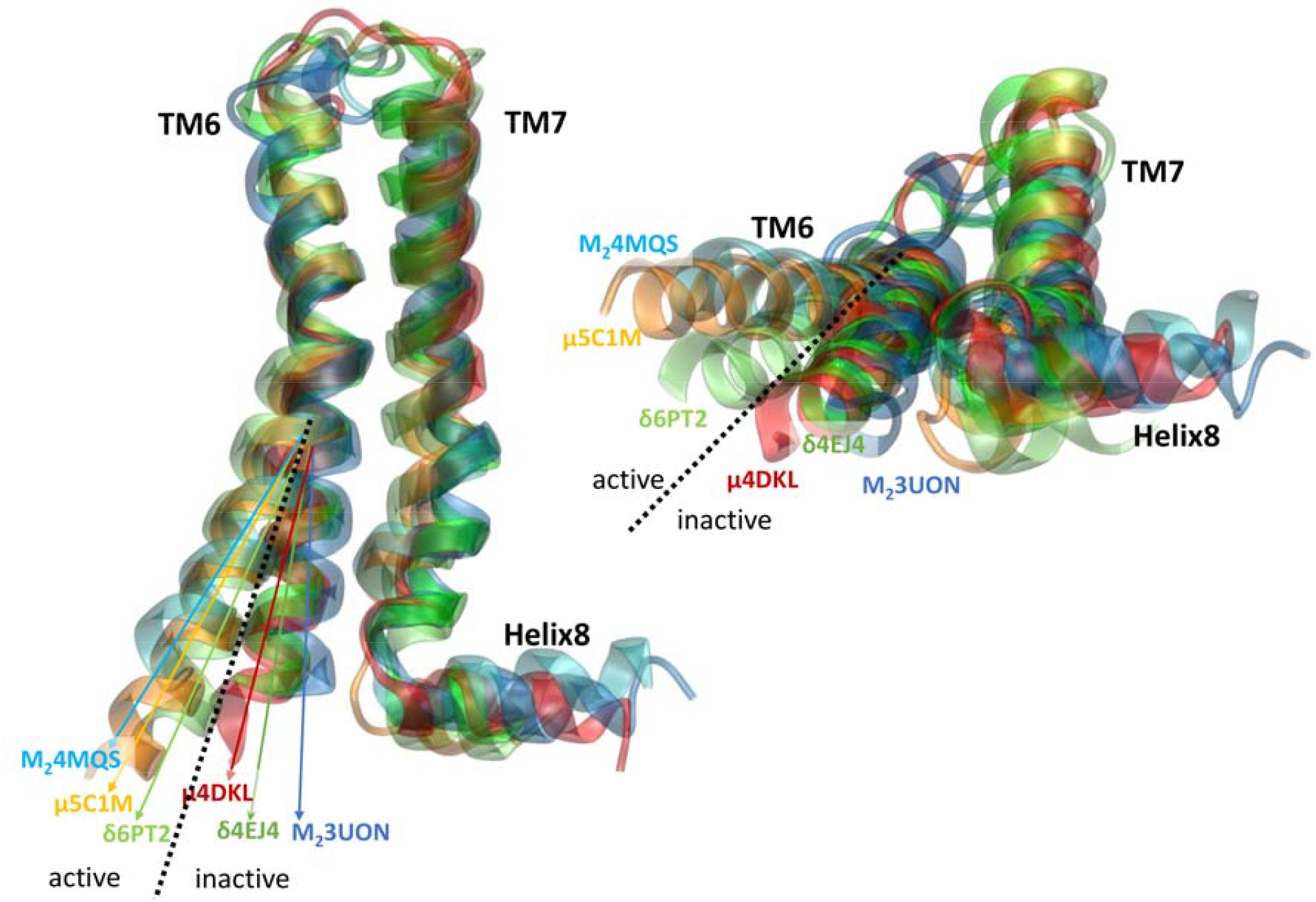
Comparison of inactive and active receptor conformations. Side (left) and intracellular view (right) of TM6, TM7 and Helix 8 of inactive M_2_-muscarinic (blue), δ- (green) and μ-opioid (red) and active M_2_-muscarinic (cyan), δ- (lime) and μ-opioid (orange) receptors.

Opioid receptors, particularly the μ- and δ-opioid subtypes, have been extensively studied with respect to their dimerisation, revealing a key role for cholesterol in promoting and stabilising dimers[14]. For the far less explored muscarinic receptors, modelling studies have suggested cholesterol-mediated oligomerisation via the TM6–TM7 interface[67]. In these models, cholesterol positions were inferred from template structures of the β_2_-adrenergic and 5-HT_2_B serotonin receptors. However, the presented study did not confirm cholesterol binding to the e6–e7 site (Table 2) likely due to the orientation of polar and acidic residues at the extracellular ends of TM6 and TM7 in muscarinic receptors, which face away from the membrane and thus cannot interact with cholesterol’s hydroxyl group. In contrast, our findings demonstrate cholesterol binding to the e6–e7 site in opioid receptors (Table 2). Such binding has been observed in crystal and cryo-EM structures of opioid receptors 4DKL, 5C1M, 6B73, and 6PT2[39, 41–43]. Notably, in the μ-opioid receptor, the e6–e7 site forms part of the dimer interface[42]. Furthermore, Zheng et al. reported that a cholesterol–palmitoyl interaction at the i7 site facilitates μ-opioid receptor homodimerisation[68].

Our findings reveal novel therapeutic opportunities by targeting non-canonical cholesterol-binding sites in GPCRs, enabling tissue-specific drug design. At muscarinic receptors, a cholesterol-binding site was shown to bind neuroactive steroids[69]. Thus, neurosteroids and steroid hormones modulating muscarinic receptors[70–72] very likely act via cholesterol-binding sites. The neurosteroid pregnanolone sulphate has been shown to increase acetylcholine release in the frontal cortex, amygdala and hippocampus and improve spatial recognition[73, 74]. The steroid whitanone blocked the negative effects of scopolamine on cholinergic signalling, exerted neuroprotective effects and promoted neuroregeneration and memory recovery[75, 76]. Like muscarinic receptors, membrane cholesterol affected the binding and function of opioid receptors[65]. Membrane cholesterol modulated the partitioning of δ-opioid receptors to membrane rafts[77]. Also, the binding of steroids to opioid receptors affecting their functional response was observed[78, 79]. A steroid capable of activating μ-opioid receptors was synthesised[80], indicating that, similar to muscarinic receptors, opioid receptors can be modulated by steroids. However, the non-genomic effects of steroids on the function of opioid receptors have not been studied in detail yet.

Various tissues differ in the content of membrane cholesterol. For example, cholesterol content is high in neurons and low in smooth muscle cells. Steroid-based pharmaceuticals, thus, may be more efficient at cholesterol lean tissues due to lesser competition with cholesterol for binding to GPCRs. Membrane cholesterol affects GPCR downstream signalling and may change the pharmacological profile of GPCR agonists[30]. Reduction of the content of membrane cholesterol attenuated signalling via δ-opioid receptors in neuronal cells but enhanced it in non-neuronal cells[81]. Therefore, in theory, natural differences in how receptors interact with membranes could enable pharmacological selectivity based on the tissue-specific composition of membranes, leading to selective effects among tissues.

In summary, this study identifies a shared “non-canonical” motif-less cholesterol-binding site in the groove between TM6 and TM7 in both muscarinic and opioid receptors, with significant implications for drug development targeting tissue-specific GPCR modulation.

## Abbreviations

CHO: Chinese ovary
GPCR: G-protein coupled receptor
GTPγS: guanosine 5’-O-[γ-thio]triphosphate
TM: transmembrane α-helix

## Acknowledgements

We thank Prof. Esam El-Fakahany, University of Minnesota and Dr. Vladimir Dolezal, Institute of Physiology CAS, for the critical reading of the manuscript.

## Author contributions

N.C. constructed mutants and performed measurements of functional responses. A.R. contributed to measurements of functional responses and data analysis. D.N. contributed to molecular modelling and measurements of functional response. J.J. conceived the experimental design, secured financing, performed molecular modelling and data analysis and finalised the manuscript. All authors contributed to the manuscript preparation and approved the final version.

## Data availability

All relevant data are contained within the manuscript.

